# Macroevolutionary changes in natural selection on codon usage reflects evolution of the tRNA pool across a budding yeast subphylum

**DOI:** 10.1101/2024.09.27.615277

**Authors:** Alexander L. Cope, Premal Shah

## Abstract

Across taxonomical domains, synonymous codons of an amino acid are found to be used at unequal frequencies within genes. This codon usage bias (CUB) is highly variable across species. Genome-wide CUB reflects a balance between adaptive and non-adaptive microevolutionary processes within a species. Variation in microevolutionary processes results in across-species variation in CUB. As CUB is tightly linked to important molecular and biophysical processes, it is critical to understand how changes to these processes are linked to changes in microevolutionary processes. We employed a population genetics model to quantify natural selection and mutation biases on a per-codon basis across the Saccharomycotina budding yeast subphylum. We found that the strength of natural selection and mutation biases varied significantly between closely related yeasts. Across-species variation in natural selection reflected the evolution of tRNA gene copy number. Additionally, we found evidence that changes to tRNA modification expression can contribute to changes in natural selection across species independent of changes to tGCN. Both lines of evidence support the link between the evolution of the tRNA pool and natural selection in codon usage through changes in the translation efficiency of a codon. Furthermore, we found that changes to tGCN often reflected changes to genome-wide GC%, suggesting changes to the tRNA pool reflect changes to mutation bias. Our work establishes how changes in microevolutionary processes impact changes in molecular mechanisms, ultimately shaping the macroevolutionary variation of a trait.

**Significance statement:** Codon usage bias (CUB) – the non-uniform usage of synonymous codons – is a feature of all genomes and varies across closely related species. Differences in CUB imply differences in the underlying microevolutionary processes (natural selection, mutation bias) driving CUB. CUB is hypothesized to be tightly linked to key molecular processes, particularly mRNA translation. We used a population genetics model to quantify natural selection and mutation bias on a per-codon basis across 327 budding yeasts. We found high variability in the microevolution of CUB and showed that changes in natural selection were correlated with the evolution of the tRNA pool. Our work establishes how variation in molecular mechanisms relates to variation in microevolution, shaping variation in a trait across species.

## Introduction

The genetic code is “degenerate.” The 61 amino acid encoding codons are translated into 20 amino acids, meaning that multiple codons must be ascribed to the same amino acid. Across all domains of life, synonymous codons are used at unequal frequencies, a phenomenon known as codon usage bias (CUB) [1–4]. The CUB in a genome reflects a balance between the microevolutionary processes of natural selection, mutation biases, and genetic drift [5, 6]. Natural selection for efficient or accurate translation (“translational selection”) is hypothesized to be the main selective pressure on synonymous mutations, with selection strongest in highly expressed genes due to their effects on the energetic burden of ribosome pausing or protein misfolding [7–10]. The translational selection hypothesis can explain the often observed correlation between codon usage and the tRNA pool, and the higher frequency of codons corresponding to more abundant tRNA in highly expressed genes [4, 8, 11, 12]. The overall CUB of a genome is primarily determined by mutation bias and drift because (1) highly expressed genes constitute only a small portion of protein-coding sequences in a genome and (2) translational selection is generally considered a weak selective pressure. Other evolutionary processes, such as GC-biased gene conversion [13, 14], also contribute to CUB within a genome and can obscure signatures of natural selection [15, 16].

CUB varies across species in terms of the degree of bias and the identity of the most frequently used synonyms [1, 4, 12, 17–21]. Variation in CUB on macroevolutionary timescales ultimately reflects variation in the underlying microevolutionary processes shaping CUB within a genome. Mechanistic evolutionary models can provide theoretically justified estimates of evolutionary parameters necessary to determine how variation in these processes leads to the observed macroevolutionary trends in CUB. Moreover, CUB is tightly linked to the molecular processes of DNA replication, repair (mutation bias), and protein synthesis (translational selection), such that macroevolutionary variation in the microevolutionary processes shaping CUB are hypothesized to reflect changes to the underlying molecular processes [2, 4, 12, 19].

Here, we used the Ribosomal Overhead Cost version of the Stochastic Evolutionary Model of Protein Production Rates (ROC-SEMPPR) [8, 22, 23] to quantify variation in natural selection and mutation biases shaping codon usage across 327 Saccharomycotina budding yeasts [12, 24]. Building on a previous examination of 49 budding yeasts [23], we find a substantial percentage (*≈* 18%) of the Saccharomycotina subphylum exhibits significant across-gene variation in codon usage not attributable to translational selection, suggestive of other non-adaptive evolutionary processes causing variation in codon usage across genes. We find multiple lines of evidence for translational selection on CUB across most species, including that shifts in natural selection correlate with changes to the tRNA pool across species. Such a correlation is often presumed, but our work directly shows how a key feature of protein synthesis (the tRNA pool) relates to natural selection on codon usage. Furthermore, we explore how mutation biases change across species, finding that they are strongly correlated with changes to GC%, but we find inconclusive support for a general role of the evolution of mismatch-repair (MMR) genes in driving changes to mutation biases (see *SI Appendix*).

## Results

We applied ROC-SEMPPR to quantify the strength and direction of natural selection and mutation bias shaping CUB across the 327 Saccharomycotina budding yeasts. ROC-SEMPPR is a population genetics model that disentangles natural selection from mutation biases by quantifying changes in codon usage as a function of gene expression (see Materials and Methods for details) [8, 22, 25]. Specifically, for a given gene *g* with gene expression *ϕ_g_*, the probability *p_g,i_* of seeing codon *i* is

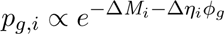

ROC-SEMPPR estimates the strength and direction of natural selection (scaled selection coefficients) Δ*η* and mutation bias Δ*M* per codon. To further clarify the meaning of these parameters, the scaled selection coefficients (from now on, just “selection coefficients”) Δ*η* ∝ *s_i,j_N_e_* in a gene of average expression, where *s_i,j_* is the unscaled selection coefficient between synonymous codons *I* and *j* and *N_e_* is the effective population size. Mutation bias 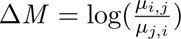 where *µ_i,j_* is the mutation rate between synonymous codons *i* and *j*. All codon-specific parameters from ROC-SEMPPR are relative to a reference codon, specifically the alphabetically last codon for each amino acid. The choice of reference codon does not affect parameter estimation. Negative values of parameter estimates indicate a codon is “favored” relative to the reference codon. Parameters are estimated using a Monte Carlo Markov Chain (MCMC) fitting procedure implemented in the **AnaCoDa** R package [26]. Previous work focused on changes in codon frequencies across the Saccharomycotina subphylum [12]. Here, we focus primarily on the variation of the selection and mutation parameters shaping these codon usage patterns.

Within-genome variation in nucleotide usage arising due to non-adaptive evolutionary processes (e.g., GC-biased gene conversion) can obscure signatures of translational selection [16, 23, 27, 28]. To account for this variation, for each species, we divided all protein-coding genes into 2 sets based on absolute codon frequencies. Specifically, we applied correspondence analysis to the absolute codon frequencies for all genes followed by CLARA clustering of the first 4 principal axes into 2 sets of genes. The genes in these two sets are potentially subject to different non-adaptive nucleotide biases. Following [23], we compared two model fits to assess the potential impact of within-genome variation in non-adaptive nucleotide biases. We refer to these models as “ConstMut” and “VarMut”. The ConstMut model assumes mutation bias Δ*M* is constant across all genes; in contrast, the VarMut model allows mutation bias Δ*M* to vary between the two sets of genes, termed the “Lower GC3 Set” and “Higher GC3 Set.” In both cases of ConstMut and VarMut models, we assume that the selection coefficients Δ*η* are the same between the two sets of genes. We then used the Spearman rank correlations between ROC-SEMPPR predicted gene expression *ϕ* and empirical gene expression determined via RNA-seq to compare our models. For this analysis, empirical estimates of gene expression were used from a subset consisting of 49 yeasts [23], with estimates for the remaining species mapped to their closest relative in that subset based on a list of one-to-one orthologs [24] (*SI Appendix*, Fig. S1). The VarMut model was considered the better model if it improved this correlation by 25% relative to the ConstMut model. The VarMut model better fit 58 of the 327 species (approximately 18%), consistent with substantial within-genome variation in non-adaptive nucleotide biases shaping the CUB of these species (*SI Appendix*, Fig. S2). For subsequent analyses, we used the selection and mutation bias estimates of the best fit of the model for each species. For simplicity, we will use the term “mutation bias” to mean any non-adaptive nucleotide bias throughout the remainder of this text, acknowledging some evolutionary processes shaping these biases are not truly mutational processes (e.g., GC-biased gene conversion).

### The selectively and mutationally favored synonyms often vary across species

To understand the evolution of codon usage on macroevolutionary timescales, it is critical to quantify the variability in the evolutionary forces shaping CUB. We determined which codons were favored by (1) natural selection and (2) mutation biases across the subphylum. Each synonymous codon was ranked from 1 (most favored) to *n* (least favored), where *n* is the number of synonyms for an amino acid, based on the selection coefficients and the mutation bias estimates for a given species.

We find that natural selection often favors the same synonym across the majority of budding yeasts under consideration; however, the degree of this bias varies across amino acids (*SI Appendix*, Fig. S3). Taking the 2-codon amino acids K-Lys, E-Glu, and Q-Gln (all NNA/NNG) as an example, the K-Lys codon AAG is generally selectively-favored relative to AAA in most species (87%), while GAA (E-Glu) and CAA (Q-Gln) are favored across 57% and 73%, respectively (Fig. 1, top left). In contrast, the NNC synonym of the 2-codon NNC/T amino acids is selectively favored in the majority of species (Fig. 1, middle left, D-Asp: 0.77%, F-Phe: 96%, H-His: 91%, N-Asn: 94%, Y-Tyr: 95%). A clear exception to this is C-Cys, with the TGT codon being selectively favored in most species (82%). The NNT or the NNC codons were selectively-favored for most species in the case of the 4-codon amino acids (including S-Ser_4_), with P-Pro generally favoring the CCA codon for most species (Fig. 1, bottom left). These results highlight that the selectively-favored codon for an amino acid often varies across the Saccharomycotina subphylum, suggesting underlying factors that alter which synonyms provide fitness advantages.

**Figure 1:**
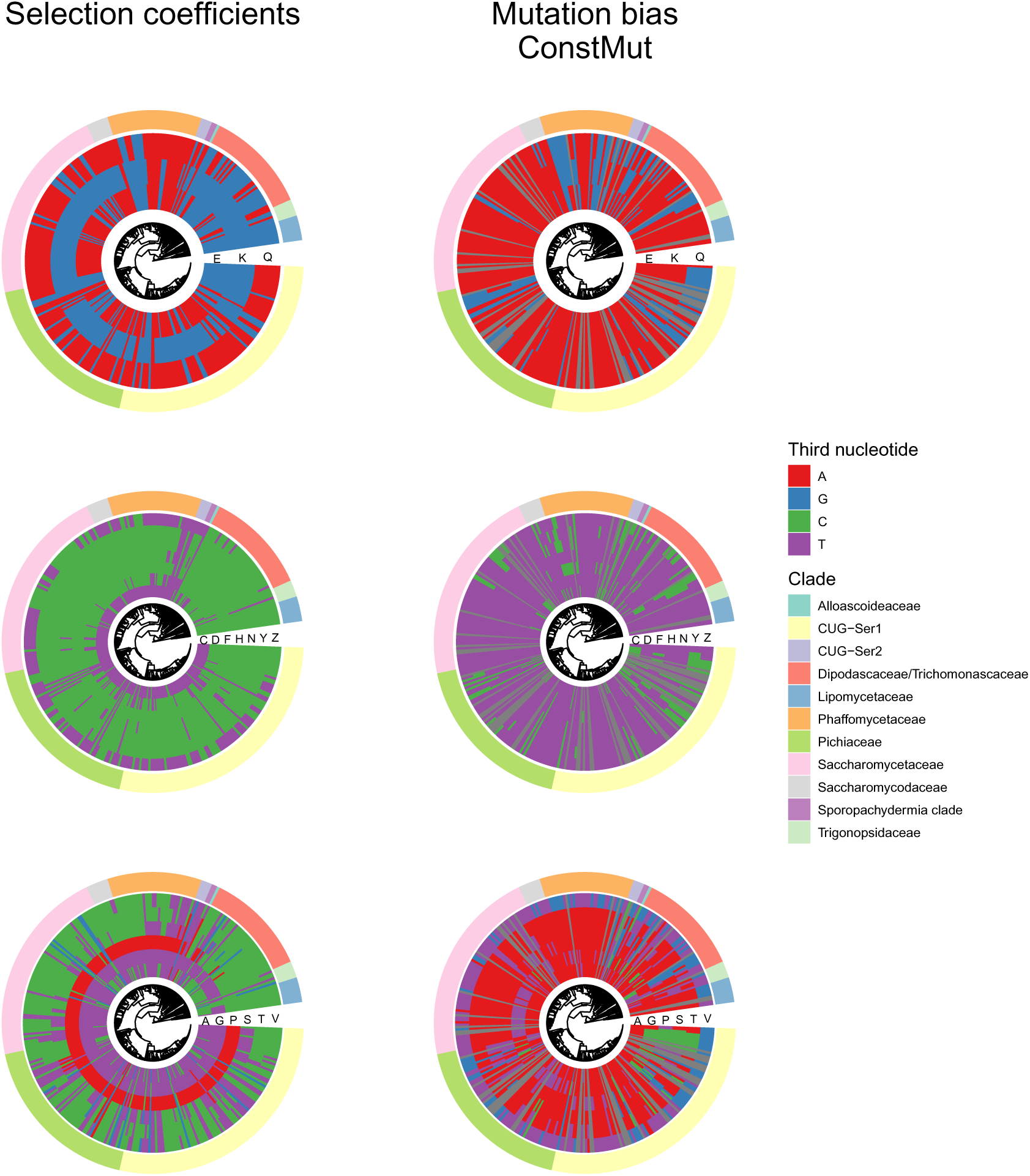
Codons favored by natural selection and mutation are largely consistent across the Saccharomycotina subphylum. Plot indicates the identity of the codon favored by natural selection and mutation bias based on estimates of (left column) selection coefficients Δη and (right column) mutation biases Δ*M*, respectively for all 2-codon and 4-codon amino acids. “Z” indicates the serine codons AGC/AGT, which are treated as separate from the other 4 serine codons. Only the ConstMut species are considered for mutation bias, so VarMut species are represented by the grey boxes. The outermost color bar indicates the major clade.

We find that mutation biases generally favor the NNA/T codons in the ConstMut species, consistent with the AT-biased genomes of the Saccharomycotina budding yeasts (Fig. 1, *SI Appendix*, Fig. S4). This pattern holds for the mutation biases of the Lower GC3% set in the VarMut species (*SI Appendix*, Fig. S5). In contrast, the mutation biases of the Higher GC3% set indicate a bias towards NNC/G synonyms (*SI Appendix*, Fig. S6). As many of these species have a genome-wide GC%*<* 50% (*SI Appendix*, Fig. S7), the evolutionary processes shaping GC% within these genes are likely to be localized to specific regions of the genome. In line with this interpretation, we find that the genome-wide GC% is consistent with estimates of mutation biases in the ConstMut and the Lower GC3% set estimated for the 4-codon amino acids. We find that the observed genomewide GC% was within the range of expected GC% based on the 6 4-codon amino acids (including S-Ser_4_) for 214 of the 327 species (*≈* 65%) (*SI Appendix*, Fig. S8). In contrast, the genome-wide GC% is consistently lower than expected based on the estimates of mutation biases from the Higher GC3% set for 51 of the 58 VarMut species (88%).

### The strength of natural selection and mutation biases shaping codon usage varies among close relatives

The above analyses indicate how the direction of natural selection and mutation biases shaping codon usage change across the Saccharomycotina subphylum. However, natural selection and mutation bias estimates not only indicate the direction of these evolutionary processes; they vary in the strength of these processes (i.e., how favored is a codon). To this end, we hierarchically clustered species based on estimates of (1) selection coefficients Δ*η* and (2) mutation biases Δ*M* to understand how these parameters varied between species and to what extent these parameters reflected shared ancestry.

The clustering of selection coefficients weakly reflects the phylogeny of the Saccharomycotina yeasts, suggesting little similarity between closely related species. While many species from the same clade fall into the same cluster, most clades are divided into separate groups (Fig. 2A, Clade color bar). We find 34 species exhibiting dramatic shifts in natural selection on codon usage relative to the remaining 293 species (Fig. 2A, row dendrogram labels 1 and 2). These species have significantly weaker correlations between ROC-SEMPPR predicted gene expression and empirical gene expression (Welch two-sample t-test *p* = 3.499E *−* 08) compared to the other 293 species, suggesting poor model fits (Fig. 2A, Emp. vs. Pred. Gene Expr. and Fig. 2B for examples from each cluster). To ensure these outlier species are not obscuring the similarity of closely related species in the hierarchical clustering, we removed them and re-performed the clustering of selection coefficients Δ*η*. We find little pairwise similarity between the clustering of selection coefficients and the phylogeny (cophenetic correlation *c* = 0.35), consistent with the notion that selection coefficients only weakly reflect the phylogeny of the Saccharomycotina yeasts.

**Figure 2:**
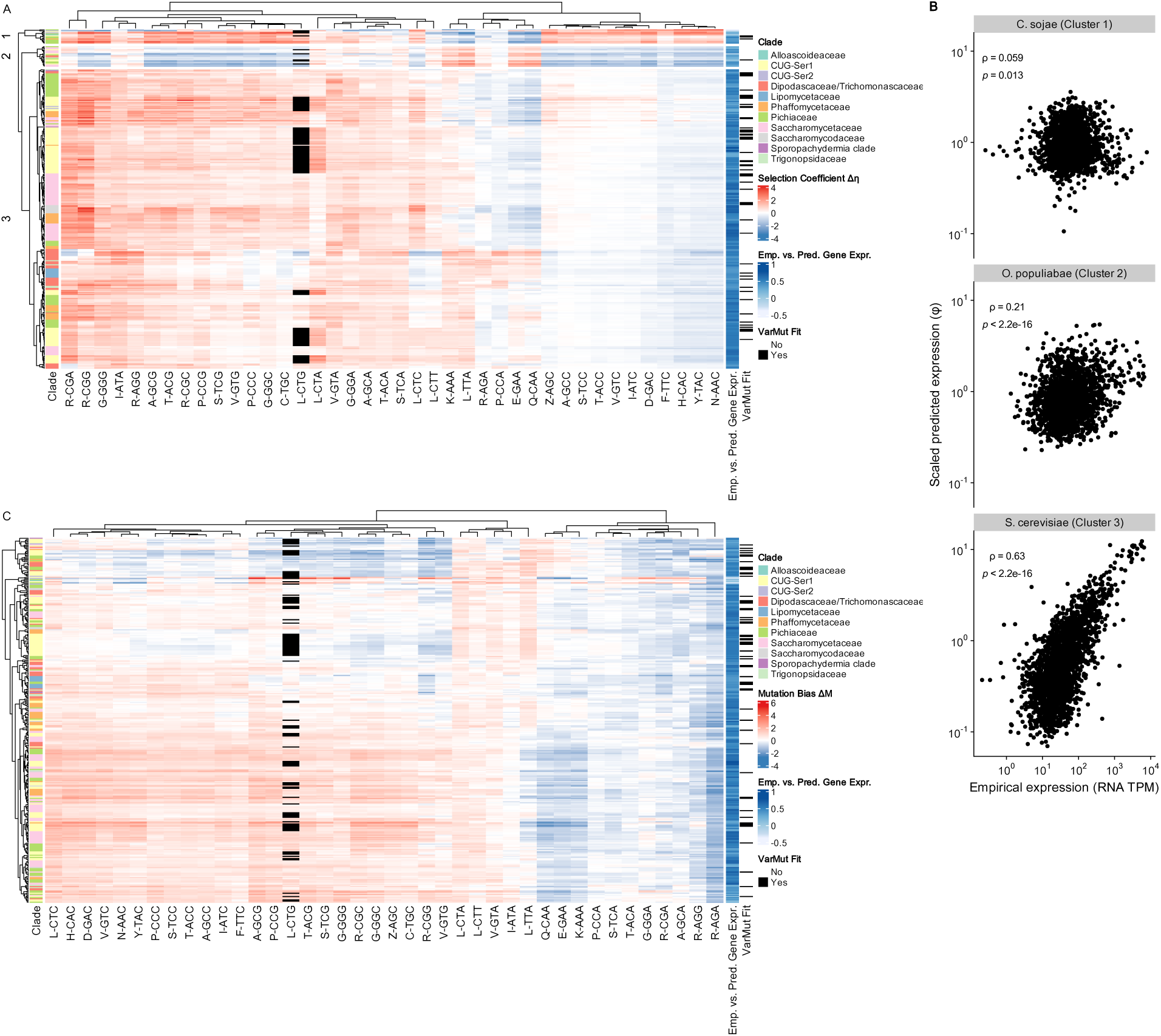
Variability of selection coefficients Δη and mutation biases Δ*M* across the Saccharomycotina subphylum. (A) Heatmap representing the selection coefficients Δη across the 327 Saccharomycotina subphylum. Red indicates a codon is selected relative to the reference synonymous codon (the alphabetically last codon for each amino acid), while blue indicates the codon is selected relative to the reference codon. Black indicates cases where the codon CTG codes for serine – in these species, CTG is treated separately from the serine codons. Row-wise dendrogram represents the hierarchical clustering of selection coefficients across species based on dissimilarity as estimated by 1 - ρ, where ρ is the Spearman correlation between the selection coefficients of two species. Numbers represent the result of splitting the clustering into 3 groups of species. Column-wise dendrogram represents the hierarchical clustering of selection coefficients across codons based on Euclidean distance. The Clade color bar indicates the major clade of the species, as defined previously[12]. The Emp. vs. Pred. Gene Expr. color bar represents the Spearman correlation between empirical gene expression and ROC-SEMPPR predicted gene expression ϕ per species. The VarMut fit color bar indicates if the model fit used for a given species was VarMut (black) or ConstMut fit (white). “Z” indicates the serine codons AGC/AGT, which are treated as separate from the other 4 serine codons. (B) Example of the correlation between empirical gene expression and ROC-SEMPPR predicted gene expression for species in the three clusters as labeled on the species-wise dendrogram in (A). (C) Same as in (A), but using mutation biases from the ConstMut and Lower GC3% set of the VarMut species. As with selection coefficients, red indicates a codon is mutationally disfavored relative to the reference codon and blue indicates it is favored.

We also examined variation in mutation biases Δ*M* across the 327 yeasts. As with the selection coefficients, we performed hierarchical clustering of mutation biases from the ConstMut species and Lower GC3% set of the VarMut species to assess their similarity between closely related species (Fig. 2, *SI Appendix*, Fig. S9 for the same analysis using ConstMut and the Higher GC3% set). Mutation biases are more variable between closely related species compared to selection coefficients as indicated by the smaller groupings of species by clade. Consistent with this qualitative observation, the cophenetic correlations between the hierarchical clustering of mutation biases and the phylogenetic tree after removing the 34 outlier species are 0.14 and 0.13, depending on the use of Lower GC3% and Higher GC3% sets for species better fit by the VarMut model.

As an orthogonal analysis to quantify the overall similarity of natural selection and mutation biases between closely related yeasts (i.e., phylogenetic signal), we estimated a multivariate version of Blomberg’s *K* (*K_multi_*) using the R package **geomorph** [29]. Selection coefficients exhibit a greater phylogenetic signal (*K_multi_* = 0.437) than mutation biases (*K_multi_* = 0.224 if using ConstMut + Lower GC3% set, *K_multi_* = 0.156 if using ConstMut + Higher GC3% set). Both analyses suggest that there exists greater variation in mutation biases between closely related species. In summary, we find that both natural selection and mutation biases are highly variable across the subphylum. The degradation of the phylogenetic signal may be due to stronger stabilizing selection on the molecular factors and processes shaping mutation bias evolution compared to those shaping natural selection (*SI Appendix*, Fig. S10).

### Selection for translation efficiency is a dominant force shaping within-genome variation in codon usage

The strength and direction of natural selection on codon usage varies across species, suggesting changes in the molecular processes shaping natural selection on codon usage. Translational selection (e.g., selection for translation efficiency/accuracy) remains the predominant adaptive hypothesis regarding genome-wide CUB, particularly in microbes. The translational selection hypothesis leads to two testable predictions: (1) codon usage will covary with gene expression and (2) highly expressed genes are biased toward codons with faster elongation rates. Such predictions are testable with ROC-SEMPPR’s estimates: (1) estimates of evolutionary-average gene expression *ϕ* are expected to be positively correlated with empirical estimates of gene expression, and (2) selection coefficients Δ*η* are expected to be positively correlated with ribosome waiting times, the inverse of elongation rates (i.e., slower codons are disfavored by natural selection).

#### ROC-SEMPPR gene expression predictions are well-correlated with empirical gene expression

After determining the best model fit between the ConstMut and VarMut models for each species, we find predicted and empirical estimates of gene expression (i.e., *ϕ* vs. RNA-seq) to be moderately to strongly correlated in most species (median Spearman rank correlation *ρ* = 0.51, with 95% of species having a correlation coefficient *>* 0.26 (Benjamini-hochberg adjusted *p <* 0.05 for all species, Fig. 3A,B). The positive correlations between predicted (i.e., based solely on codon usage) and empirical gene expression indicate prevalent translational selection on codon usage across the Saccharomycotina subphylum.

**Figure 3:**
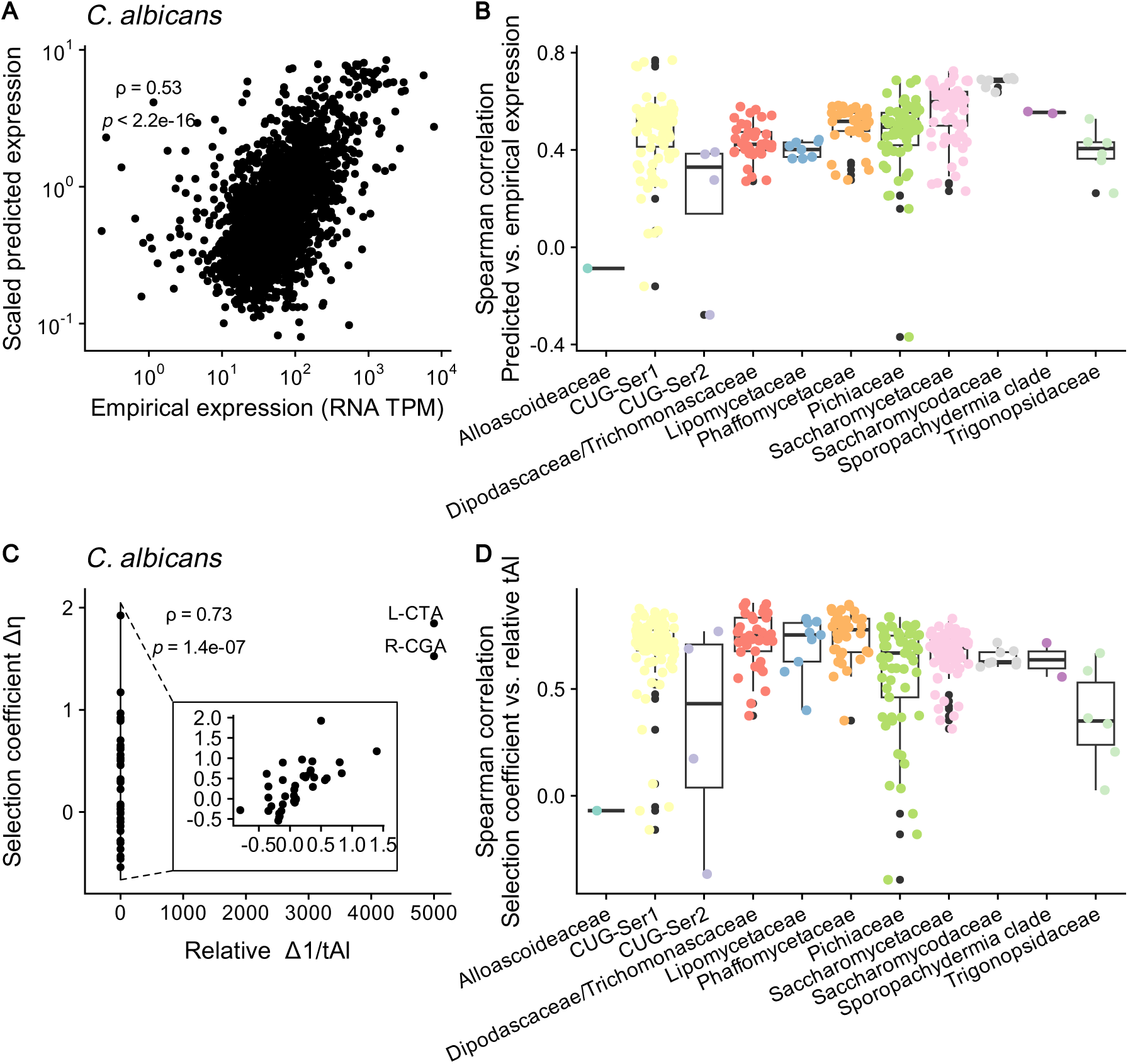
Evidence of translational selection among the Saccharomyoctina budding yeasts. (A) Example correlation between ROC-SEMPPR predicted (ϕ) and RNA-seq empirical (TPM - transcripts per million) expression for *C. albicans*. Analysis is restricted to a set of ≈ 2, 000 single-copy one-to-one orthologs across the Saccharomycotina subphylum. (B) Box plots show the distribution of Spearman rank correlations ρ computed as in (A) by clade. For most species, empirical gene expressions were taken from the closest relative for which such estimates were available. using the single-copy one-to-one orthologs. (C) Example correlation between ROC-SEMPPR selection coefficients Δη and relative tAI Δ1/tAI for *C. albicans*. Note that codons L-CTA and R-CGA are translated exclusively through wobble pairing with a large wobble penalty based on [35]. (D) Box plots showing the distribution of Spearman rank correlations ρ computed as in (C) by clade. Colored dots in (B) and (D) indicate the Spearman rank correlations for individual species and are colored by clade. Note that “Z” indicates the serine codons AGC/AGT, which are treated as separate from the other 4 serine codons.

As noted previously, a subset of 34 species exhibit weaker correlations between empirical and predicted gene expression, suggesting ROC-SEMPPR had difficulty detecting a relationship between codon usage and gene expression. This suggests two non-mutually exclusive possibilities: (1) these 34 species are subject to weak translational selection and (2) other evolutionary processes not accounted for by the VarMut model may be obscuring translational selection (i.e., model mis-specification). Possibility (1) conflicts with the observation that selection coefficients Δ*η* of these species are generally of a similar magnitude to the other 293 species (Fig. 2A). To resolve this discrepancy, we looked at underlying assumptions behind ROC-SEMPPR model fits. For instance, ROC-SEMPPR assumes gene expression to be log-normally distributed within a genome 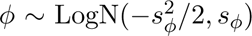, such that *E*[*ϕ*] = 1 for any species. The hyperparameter *s_ϕ_* represents the standard deviation of this lognormal distribution and is estimated along with the other parameters of ROC-SEMPPR in the **AnaCoDa** R package. Instead of shrinking selection coefficients Δ*η* towards 0 (i.e., weak selection on codon usage), ROC-SEMPPR may instead shrink *s_ϕ_* to indicate little variation in codon usage across genes. Additionally, we observe that the VarMut species generally have lower *s_ϕ_* estimates when fit by the ConstMut model, consistent with model mis-specification shrinking *s_ϕ_* (Fig. S11). Regardless, both weak translational selection and model mis-specification are expected to attenuate *s_ϕ_*, which could inflate selection coefficients. Indeed, these 34 species have a lower *s_ϕ_* on average compared to the other 293 species (Welch two-sample t-test, 0.58 vs. 0.91, *p* = 5.85E *−* 09), suggesting ROC-SEMPPR is not adequately capturing translational selection.

#### Changes in selection on codon usage are correlated with differences in tRNA gene copies between species

As selection coefficients Δ*η* reflect selection against a codon relative to its synonyms, translational selection will result in a positive correlation between selection coefficients and the expected amount of time a ribosome waits to translate a codon. In other words, if *t_i_* and *t_j_* represent the elongation rates of two synonymous codons, then

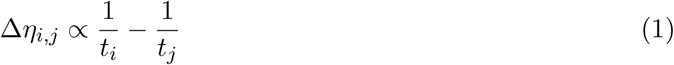

When codon *i* is slower than codon 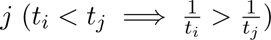, then the difference in ribosome waiting times is positive and natural selection will favor codon *j* (Δ*η >* 0). The opposite is true if codon *i* is faster than codon *j* (Δ*η <* 0). Thus, we can gain insight into the role of translational selection in shaping selection on codon usage across the Saccharomycotina subphylum by comparing our selection coefficients to molecular features known to impact ribosome waiting times. Ribosome waiting times can be estimated using high-throughput techniques such as ribosome profiling [30]; however, these measurements do not exist for the majority of the Saccharomycotina budding yeasts. Ribosome waiting times are expected to be anti-correlated with the amount of tRNA in a cell, which is itself expected to correlate with tRNA gene copy numbers (tGCNs).

Previous work found ribosome waiting times were correlated with 1/tGCNs in *S. cerevisiae* [31]. However, tRNA abundances are dynamically regulated within the cell [32, 33], potentially obscuring the relationship between tGCN and the expected ribosome waiting time of any given codon. If the tRNA pool is well-adapted to meet the translational demand of a proteome, we expect the total number of tRNA genes per amino acid to be positively correlated with amino acid usage within a species (*SI Appendix*, Fig. S12). Indeed, we find the per-amino acid tGCNs to be well-correlated with amino acid usage across the Saccharomycotina budding yeasts, with most clades having a median Spearman rank correlation *ρ >* 0.6. Based on this evidence, tGCNs appear to be a reliable, if imperfect, proxy for ribosome waiting times.

To account for reduced translation efficiency due to wobble base pairing [34], we calculated the unnormalized codon-specific weights *W* as in the tRNA adaptation index (tAI) (*SI Appendix: Supplemental Materials and Methods*, Fig. S14) [35] using code adapted from the **tAI** R package [36]. The tAI weights *W* were calculated based on the tGCNs of a given species and assuming a set of wobble rules, with codons recognized via wobble base pairing incurring a penalty reflecting a drop in translation efficiency. We used the wobble penalties originally estimated for *S. cerevisiae* [35] for all budding yeasts. Previous work showed these wobble penalties performed well in predicting protein abundances from codon usage patterns in the outgroup fungi *Schizosaccharoymces pombe* [37]. Typically, the tAI weights *W* are normalized such that they range from 0 to 1. Instead, we calculated the relative differences in the inverse of *W* i.e.,

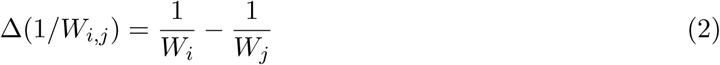

as a proxy for the relative difference in ribosome waiting times for each codon; in this case, the codon *j* is always represented by the ROC-SEMPPR designated reference codon for each of the amino acids, making Δ(1*/W_i,j_*) comparable to our selection coefficients Δ*η*. To make clear that we are using a tRNA-based proxy of waiting times, we will from here on out denote the relative tAI Δ(1*/W*) as Δ(1*/*tAI). The median Spearman rank correlation between selection coefficients and relative tAI across all 327 budding yeasts is 0.70 with an interquartile range of 0.63 to 0.76 (309/327 species *p <* 0.05, Fig. 3C,D). The positive correlation between selection coefficients and tAI indicates translational selection is a prevalent force shaping CUB across the Saccharomycotina subphylum. We observe that species showing stronger correlations between empirical and predicted gene expression tend to have stronger correlations between selection coefficients and relative tAI, but this correlation is weak (Spearman rank correlation *ρ* = 0.27, *p* = 9.9E *−* 07, *SI Appendix*, Fig. S13).

Regarding the 34 outlier species suspected of being poor model fits, these species exhibit weaker correlations between selection coefficients and relative tAI (mean Spearman rank correlation of 0.22 vs. 0.70 for the other 293 species, Welch t-test *p* = 1.613E *−* 12). This is further evidence that these 34 species were poorly fit by ROC-SEMPPR, suggestive of weak translational selection or model mis-specification. As we lacked confidence in these model fits, we excluded them from the subsequent analyses described here, leaving us with 293 species.

### Across-species variation in natural selection covaries with the evolution of the tRNA pool

The codon usage in a genome and the tRNA pool have been hypothesized to coevolve [38, 39]. Given the positive correlation between selection coefficients Δ*η* and relative tAI Δ(1*/*tAI) within species, a natural extension of this hypothesis is that changes to natural selection on codon usage across species reflect the evolution of the tRNA pool. Previous work examined the evolution of codon usage across species as it relates to the tRNA pool, but largely relied on heuristic measures [4, 12, 19, 40] or summarized selection on codon usage as a single variable [2, 27]. Consistent with previous work, we find the average strength of selection on codon usage increases with the total number of tRNA genes (Spearman rank correlation *ρ* = 0.22, *p* = 0.00025, *SI Appendix*, Fig. S15). This suggests greater investment in translation resources may lead to stronger selection for efficient translation.

#### Differences in selection coefficients of codons reflect differences in tGCN

Under translational selection, the favored codon between two synonyms is expected to have a lower waiting time i.e., when the relative tAI for two codons *i* and *j* Δ(1*/*tAI*_i,j_*) *<* 1, then selection coefficient Δ*η_i,j_ <* 1. For each codon, we examined the frequency with which selection coefficients are concordant (i.e., both positive or both negative) as relative tAI across the budding yeasts. Using Q-CAA as an example, the relative tAI and selection coefficients are concordant in 88% of species (Fig. 4A). For 32 codons of the 40 codons under consideration, selection coefficients and relative tAI are concordant for the majority of species (binomial test, Benjamini-Hochberg adjusted *p <* 0.05 for each codon), consistent with expectations under translational selection (Fig. 4B, *SI Appendix*, Fig. S16). For the other 8 codons, the direction of natural selection agreed with relative tAI in fewer species than expected by random chance (binomial test, Benjamini-Hochberg adjusted *p <* 0.05 for each codon). These 7 codons are mostly (1) codons belonging to the 4-codon amino acid groups (G-Gly, T-Thr, V-Val, P-Pro, and S-Ser_4_) and (2) being of the form NNC (all relative to a corresponding NNT codon).

**Figure 4:**
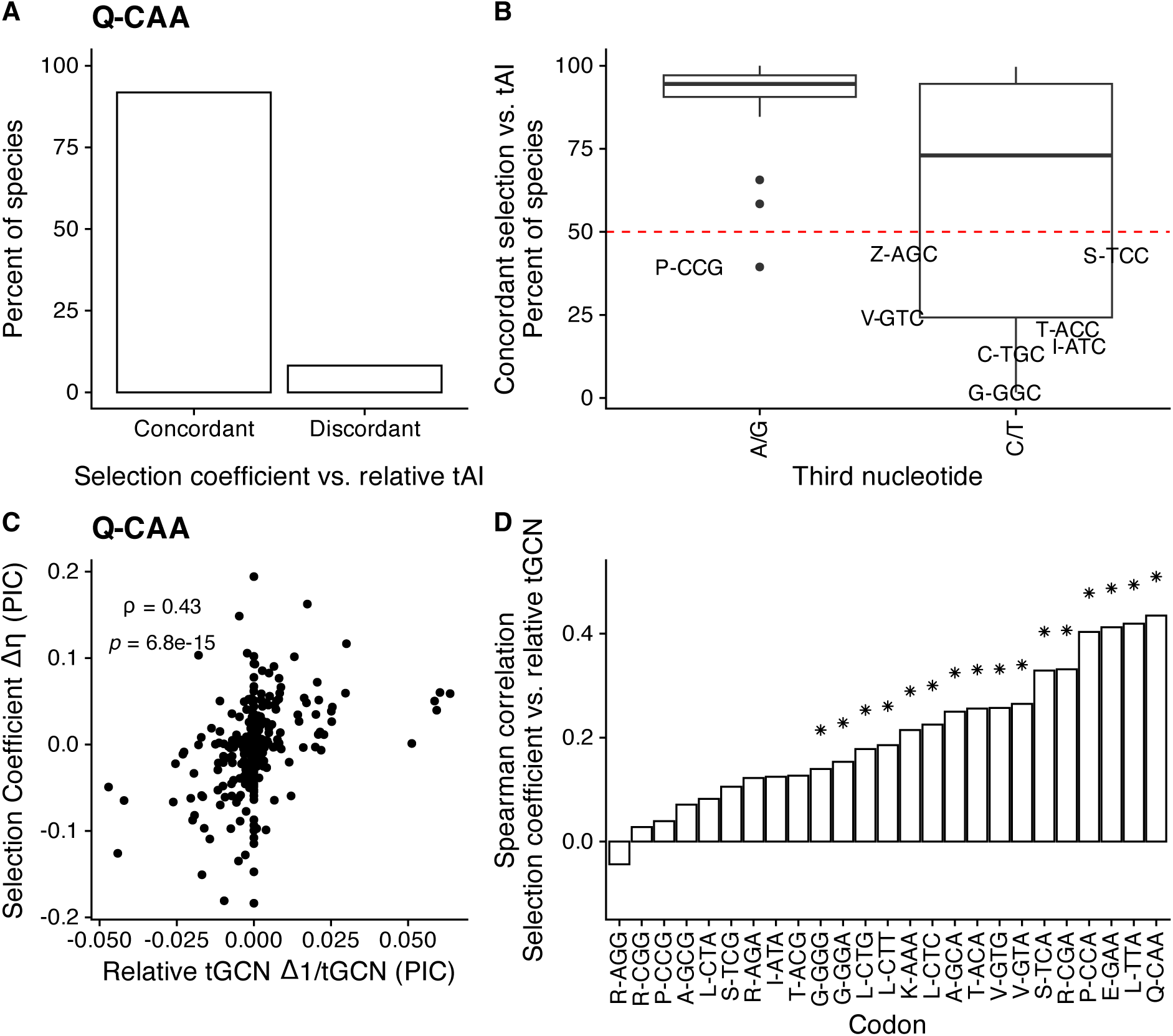
Relationship between selection coefficients and the tRNA pool across species. (A) Percent of species for which selection coefficients Δη and relative tAI Δ(1/tAI) are concordant vs. discordant for the Q-Gln codon CAA. (B) Boxplots showing the distribution for the percent of species in which selection coefficients are concordant with relative tAI. Codons are divided into either A/G-ending or C/T ending codons. Codons with a concordance percentage less than 50% are annotated (with horizontal “jitter” used to improve readability). (C) Example scatter-plot showing the relationship between selection coefficients Δη and relative tGCN Δ(1/tGCN)for Q-Gln codon CAA (relative to CAG) across species. Points represent the phylogenetic independent contrasts (PIC) of the selection coefficients and relative tGCNs. Spearman rank correlation ρ and associated p-value is reported. (D) Bar plot representing the Spearman rank correlations between selection coefficients and relative tGCN across species for all codons. “*” indicate statistical significance p < 0.05 after correcting for multiple hypothesis testing via Benjamini-Hochberg. Note that “Z” indicates the serine codons AGC/AGT, which are treated as separate from the other 4 serine codons.

The fact that selection coefficients and relative tAI generally agree does not necessarily mean changes to one variable indicate a similar change in the other. Thus, we tested if changes to natural selection correlate with the evolution of the tRNA pool. Most NNC codons exhibited little variation in the corresponding relative tAI across species, with many of these codons having fewer than 50 unique relative tAI values across the tree (reminder that NNC is relative to NNT for all 2 and 4-codon amino acids) (*SI Appendix*, Fig. S17A). This was, in part, due to NNC/T codons being read almost exclusively by the same tRNA in most species. As a result, differences in ribosome waiting time between NNC and NNT codons must be due to the effects of wobble base pairing. Consistent with this, we find NNC/T codons tend to have selection coefficients closer to 0 compared to NNA/G codons (*SI Appendix*, Fig. S17B), indicating NNC/T codons are generally less distinguishable by natural selection. As we do not allow for differences in wobble efficiency across species, we estimated the relative tGCN for across-species comparisons i.e.,

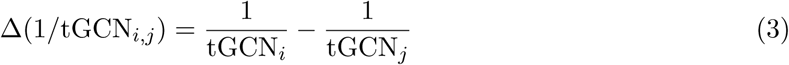

where tGCN*_i_* represents the number of tGCN that can recognize codon *i* (including through standard wobble rules). We focus on the 25 codons for which differences between synonymous codons could be ascertained based on differences in tGCN (mostly A/G-ending codons, except L-Leu codons CTC and CTT, both relative to TTG).

Of these 25 codons, 16 show a positive Spearman rank correlation between relative tGCN and selection coefficients (Fig. 4C,D, Benjamini-Hochberg adjusted *p <* 0.05). Similar results are obtained via phylogenetic generalized least squares (PGLS) that also accounted for across-species variation in GC content and *s_ϕ_* (as a potential confounder), with 15 of the 25 codons exhibiting a significant positive relationship between relative tGCN and selection coefficients (*SI Appendix*, Fig. S18). Overall, 18 of the 25 codons exhibited a statistically significant, positive relationship between selection coefficients and relative tGCN in at least one of the phylogenetic analyses. These finding support the hypothesis of coevolution of adaptive CUB and the tRNA pool.

#### Changes in selection of NAA/G codons covaries with expression of tRNA modification enzymes

Some tRNA modifications can alter the waiting times of a codon by modulating the efficiency of wobble base pairing [41, 42]. Evolutionary changes to the functionality of tRNA modification enzymes may also signal shifts in the strength or direction of natural selection on codon usage. In *S. cerevisiae*, knockouts of the multimeric Elongator Complex protein (responsible for the U34 modification mcm^5^s^2^U) resulted in increased waiting times at codons AAA, CAA, and GAA [41, 42]. Given the impact of tRNA modifications on waiting times, across-species variation in tRNA modification enzyme activity may contribute to variation in natural selection on codon usage. We examined the impact of differences in gene expression *ϕ* (as a proxy for overall enzyme activity) of the catalytic activity proteins of the Elongator Complex IKI3, ELP2, and ELP3 on changes to natural selection (see *SI Appendix: Supplemental Materials and Methods*). We observe negative correlations between predicted gene expression and selection coefficients Δ*η* for individual codons (Fig. 5A for IKI3 expression vs. AAA selection, *SI Appendix*, Fig. S19), consistent with natural selection favoring the NNA codon as expression of the modification enzymes increases. However, this did not control for other factors likely related to selection (e.g., tGCN) or expression of Elongator Complex proteins (e.g., genome-wide GC%). We performed PGLS to examine the impact of the IKI3, ELP2, and ELP3 on natural selection while controlling for changes to tGCN, genome-wide GC%, and *s_ϕ_* (Fig. 5B). Unsurprisingly, tGCN has an overall larger effect on variation in natural selection across species compared to Elongator Complex expression. However, we find that expression levels of the protein IKI3 contribute to variation in natural selection for 2 of the 3 codons. We note there is collinearity between many of our independent variables, which may result in overestimating our standard errors; however, these correlations are weak (*SI Appendix*, Fig. S20). Taken together, our results suggest changes to tRNA modification enzyme activity or expression also can reflect changes to natural selection on codon usage.

**Figure 5:**
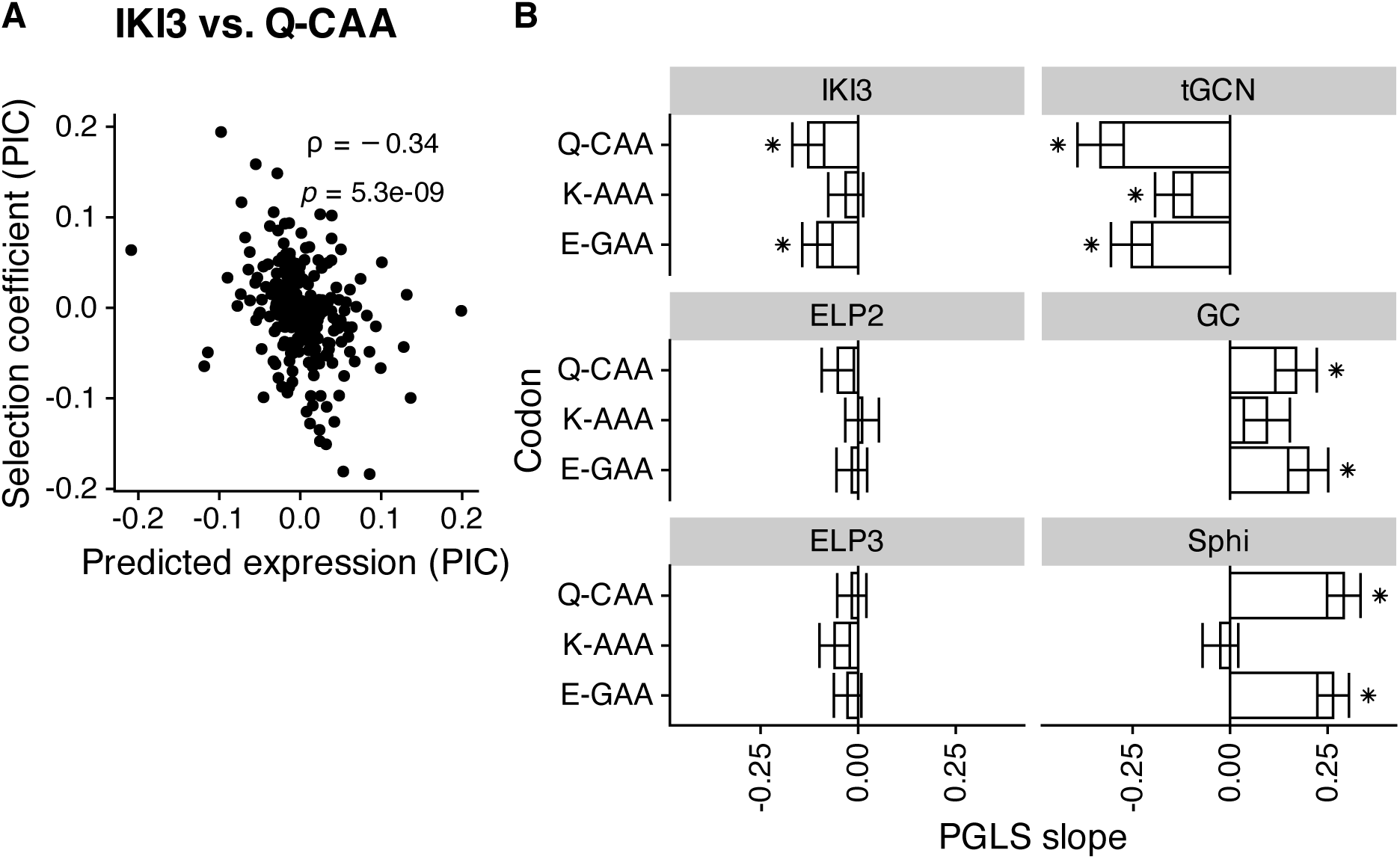
Natural selection on codon usage and covaries with gene expression of proteins forming the tRNA modification enzyme Elongator Complex. (A) Example scatter-plot showing the relationship between the selection coefficients Δη of Q-Gln codon CAA and predicted gene expression of IKI3. (B) Bar plot representing the effects (i.e., PGLS slopes) of Elongator Complex gene expression, tGCN, and genome-wide GC% on variation in selection coefficients of codons recognized by tRNA modified by the Elongator Complex. Error bars indicated ±1 std. error. “*” indicate statistical significance p < 0.05 after correcting for multiple hypothesis testing via Benjamini-Hochberg.

**Figure 6:**
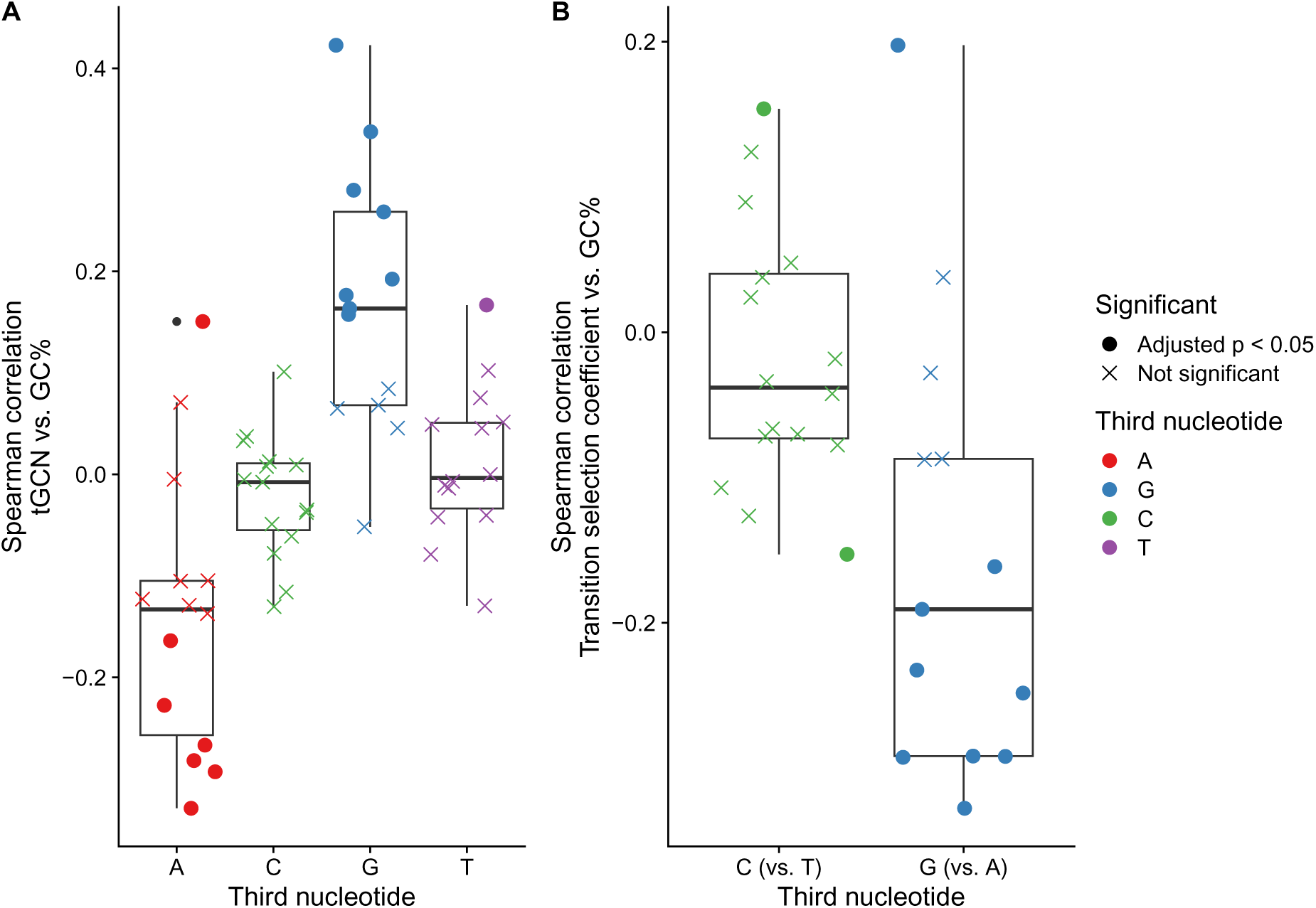
Natural selection on NNA/G codons coevolves with mutation bias. All Spearman rank correlations ρ were calculated after applying phylogenetic independent contrasts (PIC). (A) Boxplots showing distributions of Spearman rank correlations between genome-wide GC% and the absolute tGCN. (B) Boxplots showing distributions of Spearman rank correlations between genome-wide GC% and the transition selection coefficients Δη_Transition_, which reflect differences in selection between NNG and NNA codons, and NNC and NNT codons.

The greatest discrepancies between the tRNA pool and natural selection were the NNC codons of the 4-codon amino acids, which are translated almost exclusively via wobble pairing with a tRNA carrying an adenine to inosine (A to I) modification at the 34th nucleotide (I34) in most species. Variation in natural selection on these codons may reflect variation in I34 modifications. As with Elongator Complex proteins, we used the predicted gene expression values of the tRNA modification enzymes TAD2 and TAD3 as proxies for the amount of inosine modification. However, we find only one statistically significant relationship between the selection coefficients Δ*η* and the I34 modification enzyme expression (*SI Appendix*, Fig. S21). This suggests variation in natural selection of the 4-codon NNC codons (relative to the NNT synonym) is likely not due to changes in the expression of the TAD2 and TAD3 enzymes.

#### Changes in selection on some codons are due to changes in genome-wide mutation biases

Evolutionary changes to the tRNA pool may be the result of neutral evolutionary processes or an adaptation in response to changes in demand for certain tRNAs or tRNA modifications. One hypothesis is tRNA demand changes due to mutation biases [16, 43] that alter the genome-wide CUB, creating a mismatch between the codon usage of the majority of genes and the tRNA pool. This mismatch exerts a selective pressure on the tRNA pool that favors changes better aligning it with the new codon usage e.g., the gain of tRNA genes via gene duplication. This change in the tRNA pool can alter the ribosome waiting times of codons, resulting in changes to natural selection on codon usage.

Using genome-wide GC% as a general proxy for mutation biases (genome-wide GC% is strongly correlated with mutation bias Δ*M* and is well-predicted from these estimates, see *SI Appendix*, Fig. S8), we observed the absolute tGCN of NNA/G codons generally showed stronger correlations with genome-wide GC% (Wilcoxon signed rank test, *p* = 0.005 and *p* = 0.0005). The same correlations for NNC/T codons were mostly distributed around 0 (Wilcoxon signed rank test, *p* = 0.19 and *p* = 0.71). As previously noted, in the case of the 4-codon amino acids, the NNC codons are typically translated via wobble pairing through an A-to-I tRNA modification at the 34th nucleotide in the tRNA. The 2-codon amino acids encoded by NNC/T codons are almost universally read by the tRNA*_GNN_* because a tRNA*_INN_* would result in frequent missense errors. Thus, the tGCN for NNC/T codons may be less responsive to changes in GC% because of the structure of the genetic code and the inosine I34 modification.

Based on changes to the tRNA pool with GC%, we expect the selection coefficients of NNA/G codons to generally show a stronger correlation with GC% compared to NNC/T codons. As ROC-SEMPPR’s selection coefficients for each amino acid are estimated relative to a pre-defined reference (the alphabetically last codon), we calculated the “transition selection coefficients” Δ*η*_Transition_ as

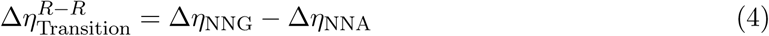

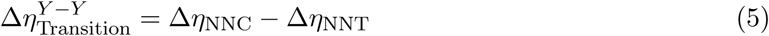

to estimate the relative strength and direction of natural selection between purines (NNA vs. NNG) and pyrimidines (NNC vs. NNT). To keep with ROC-SEMPPR’s standard interpretation of selection coefficients, Δ*η*_Transition_ *<* 0 means the NNG or NNC codon is selectively favored relative to the NNA or NNT codon, respectively. Indeed, the transition selection coefficients for NNG codons became more negative as GC% increased, consistent with stronger natural selection in favor of these codons (Wilcoxon signed rank test *p* = 0.005). We note this does not necessarily mean the NNG codons became the selectively favored codon, but simply that the selection against this codon became weaker. In contrast, the transition selection coefficients for the NNC codons were distributed around 0 (Wilcoxon signed rank test *p* = 0.25). Our results support a model in which natural selection on NNA/G codons changes in response to the evolution of the tRNA pool driven by changes to mutation biases.

## Discussion

The direction and magnitude of codon usage bias (CUB) varies across species. Theoretically justified estimates of the underlying microevolutionary processes that shape CUB within a species and how these relate to molecular mechanisms are critical for understanding the causes of the observed macroevolutionary variation in CUB. We applied a population genetics model ROC-SEMPPR to the protein-coding genes of 327 Saccharomycotina budding yeasts [12, 24] to estimate natural selection and mutation biases at codon-level resolution for each species [8, 22, 25]. The formulation of ROC-SEMPPR and its predecessors assume that only non-adaptive directional force (i.e., favoring one synonym over another) shaping codon usage bias is mutation bias, but many other relevant directional non-adaptive processes exist e.g., GC-biased gene conversion [14]. As ROC-SEMPPR quantifies natural selection on codon usage via changes to gene expression, all non-adaptive processes shaping codon usage uncorrelated with gene expression are absorbed into the mutation bias parameter. This is particularly problematic if these processes cause genome-wide variation in the non-adaptive nucleotide background: ROC-SEMPPR is most likely to mistake this variation to be the result of natural selection [23]. We built upon our recent work coupling an unsupervised machine learning approach with ROC-SEMPPR approach to identify genes subject to different non-adaptive nucleotide biases [23], finding 18% of budding yeasts exhibited significant within-genome variation in non-adaptive nucleotide biases. Previously, we found genes assigned to the different sets were largely differentiated based on GC3% and tended to be colocalized along chromosomes, leading to regions of low and high GC3% content [23]. This is suggestive of GC-biased gene conversion in regions of frequent recombination. Although we have no direct evidence of GC-biased gene conversion, Saccharomycotina yeasts better fit by the VarMut model showed clear mutation biases towards GC-ending codons in the Higher GC3% set. Further work is needed to elucidate the causes of within-genome variation in non-adaptive nucleotide biases and why these forces are more prevalent in some species of budding yeasts compared to their close relatives.

Translational selection is a prominent hypothesis for explaining adaptive CUB (i.e., CUB driven by natural selection), particularly in microbes [3]. Most of the Saccharomycotina subphylum exhibited evidence of translational selection, including moderate to strong positive correlations between predicted and empirical estimates of gene expression, and between selection coefficients Δ*η* and relative tAI (a proxy for differences in ribosome waiting times of the codons) within species. Furthermore, we found selection coefficients and relative tAI to be concordant (i.e., both positive or both negative) for the majority of species and amino acids, consistent with expectations under translational selection [5, 6, 38]. The translational selection hypothesis naturally leads to the question of why we see variation in adaptive CUB across species: variation in effective population size *N_e_* (which modulates the impact of genetic drift) or changes to unscaled selection coefficient *s* [44]? Our work focused primarily on the latter, as changes to *s* may be due to across-species changes in elongation rates. Consistent with this, we observed positive correlations between selection coefficients and relative tGCN (the difference in tGCN between codons) of the NNA/G codons. As further evidence for the role of the tRNA pool, changes to the expression levels of the Elongator Complex – a key tRNA modification enzyme – were correlated with changes to natural selection on codons AAA, CAA, and GAA across species.

Unlike with the NNA/G codons, the causes of differences in natural selection for NNC/T codons were ambiguous. For these 2-codon amino acids (except C-Cys), we still observed selection coefficients and relative tAI to be highly concordant across species, indicating the selectively favored codon is expected to be the most efficient codon. As the NNC/NNT codons of the 2-codon amino acids are recognized almost exclusively by a tRNA_GNN_, we suspect this variation in selection coefficients was primarily due to differences in the overall strength of natural selection e.g., due to differences in effective population size *N_e_* [16]. In contrast, we found clear discordance between the selection coefficients of NNC/T codons for the 4-codon amino acids, which are almost exclusively recognized by a tRNA_INN_. These results suggest (1) our calculations of relative tAI are not adequately accounting for the effects of the I34 modification on wobble efficiency or (2) other selective pressures may be shaping the codon usage of the 4-codon amino acids. Regarding possibility (1), we found no clear relationship between the C/T selection coefficients and either genome-wide GC% or inosine tRNA modification enzyme gene expression (TAD2, TAD3), both of which may serve as proxies for the expected amount of tRNA modification. One possibility is factors other than expression may impact the tRNA modification activity of TAD2 and TAD3, such as changes to protein structure. Future work will investigate how changes to tRNA modification enzyme structure relate to tRNA modification activity.

Our results provide direct evidence that across-species variation in natural selection on codon usage correlates with the evolution of the tRNA pool (i.e., due to changes in the unscaled selection coefficients *s*) at the level of individual codons. We acknowledge these correlations are weak to moderate, which may be due to several factors. First, as previously mentioned, changes in natural selection on codon usage could also be due to changes in effective population size *N_e_*. Other factors hypothesized to impact natural selection on codon usage are growth rates and optimal growth temperature [2, 16, 17, 40, 44], which are hypothesized to globally alter the unscaled selection coefficient *s*. With the current data, we cannot decompose the scaled selection coefficients Δ*η* into the effective population size and the unscaled selection coefficients. We expect these variables to dilute the across-species relationship between selection coefficients Δ*η* (∝ *sN_e_*) and the tRNA pool. Second, empirical estimates of codon-specific ribosome waiting times from ribosome profiling data are imperfectly (albeit moderately to strongly) correlated with tGCN-based proxies [45]. As regulation of tRNA abundances and gene expression can change in response to environmental stimuli, species subjected to more variable environments may have an attenuated relationship between tGCN and codon usage. Future work on this topic will benefit from the expansion of ribosome profiling into non-model systems and the emergence of sequencing technologies to more reliably quantify tRNA and tRNA modification abundances within the cell [33]. Lastly, ROC-SEMPPR averages over different selective pressures that also scale with gene expression, further obscuring the relationship between selection on codon usage and the tRNA pool. For example, selection against translation errors (missense errors, nonsense errors, etc.) is also expected to scale with gene expression [7, 46], but the most efficient codon may not always be the most accurate codon [47]. Selective pressures on CUB restricted to specific regions of a protein-coding sequence, such as selection against mRNA secondary structure around the 5’-end [48], also shape adaptive CUB. How different selective pressures interplay to shape the observed CUB remains an open question and will necessitate the development of more nuanced models that can separate different forms of adaptive CUB. Given these unaccounted for factors, we suspect our results represent a conservative estimate of the relationship between the tRNA pool and natural selection on codon usage.

Our results are consistent with the coevolution of the tRNA pool and codon usage, but what might drive the evolution of the tRNA pool? Our results support a model in which changes to mutation biases exert a selection pressure on the tRNA pool, resulting in changes to tRNA abundance or tRNA modifications that impact the direction and strength of natural selection on codon usage. A recent study examined the evolution of codon usage across 223 metazoans using methods similar in principle to our approach [16]. Despite our studies using phylogenetically distinct eukaryotes, we came to similar conclusions regarding the relationship between the tRNA pool and adaptive codon usage driven by translational selection. Noteworthy is that both studies found support for a model in which changes to mutation bias as a driver of changes to the tRNA pool, thereby shaping the strength and direction of natural selection on codon. In the case of budding yeasts, this was only the case for the NNA/G codons, whereas the tGCNs of NNC/NNT codons showed no clear relationship with genome-wide GC% (a proxy for overall mutation bias). Unlike metazoans, budding yeasts cannot incorporate queuosine into tRNA_GNN_ that can modulate the wobble efficiency of 2-codon amino acids [49], which may contribute to the lack of correlation between selection coefficients and GC%. Similar patterns relating selection on codon usage to changes in mutation bias were observed in bacteria [50]. The phylogenetic breadth of ours and previous studies suggest this may be a general model for explaining many macroevolutionary patterns in codon usage.

Perhaps our most surprising finding was that microevolutionary processes shaping CUB across the Saccharomycotina subphylum are highly variable across closely related species. This was evident by (a) the generally poor agreement between the clustering of parameters and the phylogeny and (b) the overall low estimates of phylogenetic signal based on a multivariate version of Blomberg’s *K* [29]. Interestingly, estimates of natural selection were more similar than mutation biases between closely related species. Consistent with this, estimates of stabilizing selection were generally greater for the mutation bias estimates than natural selection. This suggests the underlying molecular factors shaping mutation biases (e.g., mismatch repair) are generally under stronger stabilizing selection than those relevant to natural selection on codon usage (e.g., the tRNA pool). This should not distract from the larger point that both traits are highly variable across species, ultimately driving variation in CUB.

Hierarchical clustering revealed that 34 of the 327 budding yeast were poorly fit by ROC-SEMPPR, at least relative to other species. On average, ROC-SEMPPR parameter estimates for these species were more weakly correlated with empirical estimates of gene expression and codon-specific waiting times. These correlations were often positive, suggesting possible isolated shifts in the direction of natural selection acting on codon usage. Based on the poor model fit, it is possible that these species (1) have reduced natural selection acting on codon usage below the drift barrier [51], possibly due to reduced effective population sizes or (2) additional evolutionary forces acting on codon usage that further obscure signals of natural selection related to protein synthesis. These species serve as excellent starting points for future studies to elucidate the complex interplay of evolutionary forces that shape CUB.

Our work demonstrates the utility of applying a population genetics approach in a phylogenetic context. The explicit estimation of population genetics parameters provided insights into how a microevolutionary process (natural selection) changed over macroevolutionary time scales. In doing so, we were able to link these changes in natural selection to changes in the evolution of a molecular feature (the tRNA pool), suggesting a link between evolution and a molecular process (peptide elongation). Improvements in data availability, such as ribosome profiling data for more species, and population genetics models that explicitly incorporate other molecular processes, such as biased gene conversion or other selective pressures, will provide deeper insights into the factors shaping the evolution of codon usage within and across species.

## Materials and methods

We obtained genome sequences, associated annotation files, the Saccharomycotina species tree, and a list of orthologs from previous work [24]. We excluded mitochondrial genes and protein-coding genes with non-canonical start codons, internal stop codons, and sequences whose lengths were not a multiple of three from all analyses. We queried all protein sequences against a BLAST database built from sequences in the MiToFun database to identify and remove mitochondrial sequences (http://mitofun.biol.uoa.gr/).

Empirical gene expression measurements were taken from [23] and the sources cited therein. Briefly, adapters for each sequence were trimmed using **fastp** [52], and genes were quantified using **kallisto** [53]. Transcripts-per-million (TPMs) were re-calculated for each transcript by rounding raw read counts to the nearest whole number [54].

We ran the latest version of tRNA-ScanSE [55] using the default settings for eukaryotic genomes on the genome FASTA files for each yeast with mitochondrial genomes removed. This provided us with predicted tRNA genes for each species, allowing us to calculate the tRNA gene copy numbers (tGCNs) for each species under consideration. tRNA genes identified as “pseudo” genes were ignored when calculating the tGCNs.

Statistical analyses were peformed using the R programming language (v4.4.0). Multiple hypothesis testing was accounted for via the Benjamini-Hochberg procedure with a false discovery rate of 0.05.

### Analyzing codon usage patterns with ROC-SEMPPR

The Ribosomal Overhead Cost version of the Stochastic Evolutionary Model of Protein Production Rates (ROC-SEMPPR) is implemented in a Bayesian framework. This allows for simultaneous estimation of codon-specific selection coefficients and mutation bias, as well as gene-specific estimates of evolutionary average gene expression by assuming that gene expression follows a log-normal distribution [22]. ROC-SEMPPR does not require empirical gene expression data, meaning it can be applied to any species with annotated protein-coding genes.

ROC-SEMPPR is formulated in terms of the expected cost-benefit ratio 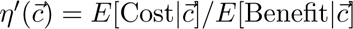 of a gene using a sequence of synonymous codons 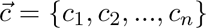 for a protein-coding sequence of length *n*. The “cost” represents the overhead cost of ribosome pausing in terms of NTP during mRNA translation, with less efficient synonymous codons resulting in higher costs. For tractability, we assume 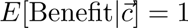, such that all differences in 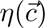 are due to differences in 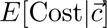. To represent the overhead cost of ribosome pausing, the 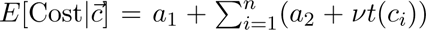. Here, *a*_1_ and *a*_2_ are constants representing the amount of NTP expended on translation initiation and peptide elongation, respectively. The function *t*(*c_i_*) represents the average waiting or pausing time a ribosome spends at codon *c_i_* a *ν* is a constant scaler that converts this pausing time to units of NTP. If we treat *ϕ_g_* as the target gene expression (specifically, the protein production rate) of gene *g*, then 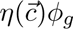 represents the average energy flux an organism must expend to meet its target production rate for a given gene. If we assume that every NTP spent per unit time leads to a small, proportional reduction in genotype fitness, then the fitness of a given coding sequence is

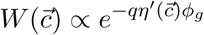

where *q* is a scaler representing the reduction in fitness per NTP expended.

Assuming independence between sites, such as when natural selection on codon usage is to minimize the cost of protein synthesis due to ribosome pausing, the expected frequencies of codons for an amino acid *aa* in gene *g* can be modeled as a multinomial distribution. Thus, for any amino acid with *n_aa_* synonymous codons, the probability *p_i,g_* of observing codon *i* in gene *g* is defined by the equation

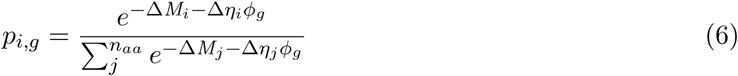

where Δ*M_i_* and Δ*η_i_* represent mutation bias and the scaled selection coefficient (i.e. 2*N_e_s*) of codon *i* relative to a reference synonymous codon (arbitrarily chosen as the alphabetically last codon), and *ϕ_g_* represents gene expression of gene *g* which follows from the steady-state distribution of fixed genotypes under selection-mutation-drift equilibrium [8, 22, 56]. Specifically, 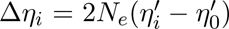, where *N_e_* is the effective population size and 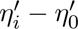 represents the difference in the expected cost of a coding sequence if using synonymous codon *i* instead of the reference synonym (i.e., *i* = 0) for a given amino acid. The mutation bias parameter Δ*M_i_* represents the log of the ratio of mutation rates between the synonymous codon *i* and the reference synonym, i.e. 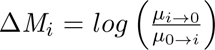, where *µ_i→j_* represents the mutation rate from codon *i* to *j*. Importantly, this formulation based on relative differences between synonymous codons allows the composite parameters to be estimated, such that the exact values of other key parameters (e.g., *N_e_*) are unnecessary to know or estimate directly.

For each gene, the observed codon counts for an amino acid are expected to follow a multinomial distribution with the probability of observing a codon defined by Equation 6. Given the codon counts and the assumption that gene expression *ϕ* follows a lognormal distribution 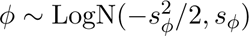 such that *E*[*ϕ*] = 1, ROC-SEMPPR estimates the parameters (including *s_ϕ_*) best fitting the codon counts via a Markov Chain Monte Carlo simulation approach (MCMC). The posterior means for the relevant parameters were used for subsequent analyses both within and across species.

ROC-SEMPPR was fit to 327 species using the R package AnaCoDa [26]. For each species, the MCMC chains were run for 200,000 iterations, keeping every 10^th^ iteration. The first 50,000 iterations were discarded as burn-in. Two separate MCMC chains were run for each species and parameter estimates were compared to assess convergence. Previous work with ROC-SEMPPR separated serine codons TCN (where N is any of the other four nucleotides) and AGY (where Y is C or T) into separate groups of codons for the analysis [8, 22]. ROC-SEMPPR assumes each mutation introduced to a population is fixed or lost before the arrival of the next mutation (i.e., “weak mutation” *N_e_µ <<* 1). The model also assumes fixed amino acid sequences for all genes. As a result, going between these two groups of serine codons would require the fixation of a non-serine amino acid before returning to serine via the fixation of another mutation, violating the fixed amino acid sequence assumption. A local version of AnaCoDa was created to handle species for which CTG codes for serine. For these species, CTG was treated as a third codon group for serine, similar to ATG (methionine) or TGG (tryptophan), which have no synonyms.

### Identifying within-genome variation in codon usage bias

We recently found numerous Saccharomycotina yeasts exhibit variation in the non-adaptive nucleotide biases shaping GC% within a genome that obscures signals of natural selection on codon usage [23]. We followed the same procedure to hypothesize genes evolving under different non-adaptive nucleotide biases across all 327 budding yeasts. For each species, correspondence analysis (CA) was applied to the absolute codon frequencies of each annotated protein-coding gene using the **ca** R package [57]. Genes were clustered into two groups based on the first four principal components from the CA using the CLARA algorithm implemented in the **cluster** R package, which is designed to perform k-medoids clustering on large datasets [58]. See our previous work for more details on the CLARA clustering algorithm [23].

ROC-SEMPPR predictions of gene expression *ϕ* for each protein-coding sequence were compared to empirical estimates of mRNA abundance using the Spearman correlation coefficient using processed data from our previous work [23]. For species lacking RNA-seq data, we compared each species’ predicted gene expression to the empirical gene expression of its closest relative for which the latter was available. This is reasonable given that mRNA abundances in yeasts evolve under stabilizing selection [59]. The VarMut model was considered an improved fit over the ConstMut model if the correlation between predicted and empirical gene expression estimates was 25% greater relative to the ConstMut model.

### Comparing codon-specific parameters across species

Across-species and across-codon variation in selection coefficients Δ*η* and mutation bias Δ*M* were compared using hierarchical clustering using the “complete linkage” algorithm with distances determined by the 1 *− ρ*, where *ρ* is the pairwise Spearman rank correlation. Results were visualized using heatmaps as implemented in the R package **ComplexHeatmap**. For each codon-specific parameter estimate, we quantified the similarity between the phylogenetic tree and the hierarchical clustering via a cophenetic correlation, which measures how well two dendrograms preserve the pairwise distances between data points. We also quantified the overall phylogenetic signal (i.e., how similar species are to their closest relatives) via a multivariate version of Blomberg’s K (*K_multi_*) as implemented in the R package **geomorph**.

Across-species correlations between traits were assessed after performing phylogenetic independent contrasts (PIC) as implemented in the R package **ape**. Phylogenetic regressions were performed using the R package **phylolm** using Pagel’s *λ* model [60].

## Acknowledgments

We thank Michael Gilchrist, Edward Wallace, Matthew Pennell, Abigail LaBella, and Antonis Rokas for their helpful comments and discussions throughout this project. A.L.C. was supported by the NIH-funded Rutgers INSPIRE IRACDA Postdoctoral Program (grant #GM093854). Additionally, this work was supported by the National Science Foundation (DBI 1936046 to P.S.) and the National Institutes of Health (R35 GM124976 to P.S.).

## Competing interests

P.S. is a Director at a stealth-mode biotech.

## Data availability

All data are publicly available via the citations provided in [12, 23, 24]. Scripts and R notebooks for re-creating our analysis and visualizations can be found at https://github.com/acope3/Saccharomycotina_subphylum_analysis.

## Notes

https://github.com/acope3/Saccharomycotina_subphylum_analysis

### Summary of Updates

This is a heavily revised version of the manuscript with updated analyses. The tAI weights are now calculated using the original wobble penalties estimated for S. cerevisiae in yeast. We examine how tGCN changes with GC content. We also now look at how often the favored codon varies across species.

## References

[1] Knight RD, Freeland SJ, Landweber LF. A simple model based on mutation and selection explains trends in codon and amino-acid usage and GC composition within and across genomes. Genome biology. 2001 4;2:research0010.1. Available from: https://genomebiology.biomedcentral.com/articles/10.1186/gb-2001-2-4-research0010.

[2] Sharp PM, Bailes E, Grocock RJ, Peden JF, Sockett RE. Variation in the strength of selected codon usage bias among bacteria. Nucleic Acids Research. 2005;33:1141–53.

[3] Plotkin JB, Kudla G. Synonymous but not the same: the causes and consequences of codon bias. Nature Reviews Genetics. 2011;12:32–42.

[4] Novoa EM, Pavon-Eternod M, Pan T, Pouplana LRD. A role for tRNA modifications in genome structure and codon usage. Cell. 2012 3;149:202–13. Available from: http://www.cell.com/article/S0092867412002127/fulltexthttp://www.cell.com/article/S0092867412002127/abstracthttps://www.cell.com/cell/abstract/S0092-8674(12)00212-7.

[5] Li WH. Models of nearly neutral mutations with particular implications for nonrandom usage of synonymous codons. Journal of Molecular Evolution. 1987 4;24:337–45. Available from: https://link.springer.com/article/10.1007/BF02134132.

[6] Bulmer M. The selection-mutation-drift theory of synonymous codon usage. Genetics. 1991;129:897–907.

[7] Drummond DA, Wilke CO. Mistranslation-Induced Protein Misfolding as a Dominant Constraint on Coding-Sequence Evolution. Cell. 2008;134:341–52.

[8] Shah P, Gilchrist M. Explaining complex codon usage patterns with selection for translational efficiency, mutation bias, and genetic drift. Proceedings of the National Academy of Sciences of the United States of America. 2011;108:10231–6.

[9] Shah P, Ding Y, Niemczyk M, Kudla G, Plotkin JB. Rate-Limiting Steps in Yeast Protein Translation. Cell. 2013;153:1589–601.

[10] Frumkin I, Lajoie MJ, Gregg CJ, Hornung G, Church GM, Pilpel Y. Codon usage of highly expressed genes affects proteome-wide translation efficiency. Proceedings of the National Academy of Sciences of the United States of America. 2018 5;115:E4940–9.

[11] Ikemura T. Correlation between the abundance of Escherichia coli transfer RNAs and the occurrence of the respective codons in its protein genes: A proposal for a synonymous codon choice that is optimal for the E. coli translational system. Journal of Molecular Biology. 1981 9;151:389–409. Available from: https://linkinghub.elsevier.com/retrieve/pii/0022283681900036.

[12] Labella AL, Opulente DA, Steenwyk JL, Hittinger CT, Rokas A. Variation and selection on codon usage bias across an entire subphylum. PLoS Genetics. 2019 7;15:e1008304. Available from: 10.1371/journal.pgen.1008304.

[13] Duret L, Galtier N. Biased gene conversion and the evolution of mammalian genomic landscapes. Annual Review of Genomics and Human Genetics. 2009 9;10:285–311. Available from: www.annualreviews.org.

[14] Galtier N, Roux C, Rousselle M, Romiguier J, Figuet E, Gĺemin S, et al. Codon Usage Bias in Animals: Disentangling the Effects of Natural Selection, Effective Population Size, and GC-Biased Gene Conversion. Molecular Biology and Evolution. 2018 5;35:1092–103. Available from: https://academic.oup.com/mbe/article/35/5/1092/4829954.

[15] de Oliveira JL, Morales AC, Hurst LD, Urrutia AO, Thompson CRL, Wolf JB. Inferring Adaptive Codon Preference to Understand Sources of Selection Shaping Codon Usage Bias. Molecular Biology and Evolution. 2021 7;38:3247–66. Available from: https://academic.oup.com/mbe/article/38/8/3247/6237909.

[16] Bènitière F, Lefébure T, Duret L. Variation in the fitness impact of translationally optimal codons among animals. Genome Research. 2025 3;35:446–58. Available from: https://genome.cshlp.org/content/early/2025/02/10/gr.279837.124.

[17] Botzman M, Margalit H. Variation in global codon usage bias among prokaryotic organisms is associated with their lifestyles. Genome Biology. 2011 10;12:R109. Available from: https://link.springer.com/articles/10.1186/gb-2011-12-10-r109https://link.springer.com/article/10.1186/gb-2011-12-10-r109.

[18] Clément Y, Sarah G, Holtz Y, Homa F, Pointet S, Contreras S, et al. Evolutionary forces affecting synonymous variations in plant genomes. PLOS Genetics. 2017 5;13:e1006799. Available from: https://journals.plos.org/plosgenetics/article?id=10.1371/journal.pgen.1006799.

[19] Wint R, Salamov A, Grigoriev IV. Kingdom-Wide Analysis of Fungal Protein-Coding and tRNA Genes Reveals Conserved Patterns of Adaptive Evolution. Molecular Biology and Evolution. 2022 2;39. Available from: https://academic.oup.com/mbe/article/39/2/msab372/6513383.

[20] Weibel CA, Wheeler AL, James JE, Willis SM, Masel J. A new codon adaptation metric predicts vertebrate body size and tendency to protein disorder. eLife. 2023;12:RP87335. Available from: https://elifesciences.org/reviewed-preprints/87335.

[21] Kotari I, Kosiol C, Borges R. The Patterns of Codon Usage between Chordates and Arthropods are Different but Co-evolving with Mutational Biases. Molecular Biology and Evolution. 2024 5;41:msae080. Available from: 10.1093/molbev/msae080.

[22] Gilchrist MA, Chen WC, Shah P, Landerer CL, Zaretzki R. Estimating Gene Expression and Codon-Specific Translational Efficiencies, Mutation Biases, and Selection Coefficients from Genomic Data Alone. Genome Biology and Evolution. 2015;7:1559–79.

[23] Cope AL, Shah P. Intragenomic variation in non-adaptive nucleotide biases causes underestimation of selection on synonymous codon usage. PLOS Genetics. 2022 6;18:e1010256.

[24] Shen XX, Opulente DA, Kominek J, Zhou X, Steenwyk JL, Buh KV, et al. Tempo and Mode of Genome Evolution in the Budding Yeast Subphylum. Cell. 2018 11;175:1533–45.e20.

[25] Wallace EWJ, Airoldi EM, Drummond DA. Estimating Selection on Synonymus Codon Usage from Noisy Experimental Data. Molecular Biology and Evolution. 2013;30:1438–53.

[26] Landerer C, Cope A, Zaretzki R, Gilchrist MA. AnaCoDa: analyzing codon data with Bayesian mixture models. Bioinformatics. 2018;34:bty138. Available from: 10.1093/bioinformatics/bty138.

[27] dos Reis M, Wernisch L. Estimating Translational Selection in Eukaryotic Genomes. Molecular Biology and Evolution. 2009 2;26:461. Available from: /pmc/articles/PMC2639113//pmc/articles/PMC2639113/?report=abstract https://www.ncbi.nlm.nih.gov/pmc/articles/PMC2639113/.

[28] Landerer C, O’Meara BC, Zaretzki R, Gilchrist MA. Unlocking a signal of introgression from codons in Lachancea kluyveri using a mutation-selection model. BMC Evolutionary Biology. 2020 8;20:109. Available from: https://bmcevolbiol.biomedcentral.com/articles/10.1186/s12862-020-01649-w.

[29] Adams DC. A Generalized K Statistic for Estimating Phylogenetic Signal from Shape and Other High-Dimensional Multivariate Data. Systematic Biology. 2014 9;63:685–97. Available from: 10.1093/sysbio/syu030.

[30] Ingolia NT, Ghaemmaghami S, Newman JRS, Weissman JS. Genome-wide analysis in vivo of translation with nucleotide resolution using ribosome profiling. Science. 2009 4;324:218–23.

[31] Wu Q, Medina SG, Kushawah G, Devore ML, Castellano LA, Hand JM, et al. Translation affects mRNA stability in a codon-dependent manner in human cells. eLife. 2019 4;8.

[32] Torrent M, Chalancon G, Groot NSD, Wuster A, Babu MM. Cells alter their tRNA abundance to selectively regulate protein synthesis during stress conditions. Science Signaling. 2018 9;11. Available from: https://stke.sciencemag.org/content/11/546/eaat6409https://stke.sciencemag.org/content/11/546/eaat6409.abstract.

[33] Behrens A, Rodschinka G, Nedialkova DD. High-resolution quantitative profiling of tRNA abundance and modification status in eukaryotes by mim-tRNAseq. Molecular Cell. 2021 4;81:1802–15.e7.

[34] Yarus M, Valle M, Frank J. A twisted tRNA intermediate sets the threshold for decoding. RNA. 2003;9:384–5.

[35] dos Reis M, Savva R, Wernisch L. Solving the riddle of codon usage preferences: a test for translational selection. Nucleic Acids Research. 2004;32:5036–44.

[36] dos Reis M. tAI: The tRNA adaptation index; 2016. R package version 0.2.

[37] Sabi R, Tuller T. Modelling the efficiency of codon-tRNA interactions based on codon usage bias. DNA Research. 2014 10;21:511–26.

[38] Bulmer M. Coevolution of codon usage and transfer RNA abundance. Nature. 1987;325:728–30.

[39] Higgs PG, Ran W. Coevolution of Codon Usage and tRNA Genes Leads to Alternative Stable States of Biased Codon Usage. Molecular Biology and Evolution. 2008 11;25:2279–91. Available from: 10.1093/molbev/msn173.

[40] Vieira-Silva S, Rocha EPC. The Systemic Imprint of Growth and Its Uses in Ecological (Meta)Genomics. PLoS Genetics. 2010;6:1–15.

[41] Nedialkova DD, Leidel SA. Optimization of Codon Translation Rates via tRNA Modifications Maintains Proteome Integrity. Cell. 2015 6;161:1606–18.

[42] Chou HJ, Donnard E, Gustafsson HT, Garber M, Rando OJ. Transcriptome-wide Analysis of Roles for tRNA Modifications in Translational Regulation. Molecular Cell. 2017 12;68:978–92.e4.

[43] Hershberg R, Petrov DA. Selection on Codon Bias. Annual Review of Genetics. 2008 12;42:287–99. Available from: http://www.annualreviews.org/doi/10.1146/annurev.genet.42.110807.091442.

[44] Subramanian S. Nearly Neutrality and the Evolution of Codon Usage Bias in Eukaryotic Genomes. Genetics. 2008 4;178:2429–32. Available from: 10.1534/genetics.107.086405.

[45] Wu CCC, Zinshteyn B, Wehner KA, Green R. High-Resolution Ribosome Profiling Defines Discrete Ribosome Elongation States and Translational Regulation during Cellular Stress. Molecular Cell. 2019 3;73:959–70.e5.

[46] Kurland CG. Translational Accuracy and the Fitness of Bacteria. Annual Review of Genetics. 1992 12;26:29–50. Available from: http://www.annualreviews.org/doi/10.1146/annurev.ge.26.120192.000333.

[47] Shah P, Gilchrist MA. Effect of Correlated tRNA Abundances on Translation Errors and Evolution of Codon Usage Bias. PLoS Genetics. 2010 9;6:e1001128.

[48] Hockenberry AJ, Sirer MI, Amaral LAN, Jewett MC. Quantifying Position-Dependent Codon Usage Bias. Molecular Biology and Evolution. 2014;31:1880–93.

[49] Zallot R, Brochier-Armanet C, Gaston KW, Forouhar F, Limbach PA, Hunt JF, et al. Plant, animal, and fungal micronutrient queuosine is salvaged by members of the DUF2419 protein family. ACS Chemical Biology. 2014 8;9:1812–25. Available from: https://pubs.acs.org/doi/full/10.1021/cb500278k.

[50] Hershberg R, Petrov DA. General Rules for Optimal Codon Choice. PLOS Genetics. 2009 7;5:e1000556. Available from: https://journals.plos.org/plosgenetics/article?id= 10.1371/journal.pgen.1000556.

[51] Lynch M. The frailty of adaptive hypotheses for the origins of organismal complexity. Proceedings of the National Academy of Sciences of the United States of America. 2007 5;104:8597–604. Available from: www.nasonline.org/adaptationandcomplexdesign.

[52] Chen S, Zhou Y, Chen Y, Gu J. Fastp: An ultra-fast all-in-one FASTQ preprocessor. Bioinformatics. 2018 9;34:i884–90. Available from: https://github.com/OpenGene/fastp.

[53] Bray NL, Pimentel H, Melsted P, Pachter L. Near-optimal probabilistic RNA-seq quantification. Nature Biotechnology. 2016 5;34:525–7. Available from: http://www.nature.com/.

[54] Wagner GP, Kin K, Lynch VJ. Measurement of mRNA abundance using RNA-seq data: RPKM measure is inconsistent among samples. Theory in Biosciences. 2012 12;131:281–5. Available from: http://link.springer.com/10.1007/s12064-012-0162-3.

[55] Chan PP, Lin BY, Mak AJ, Lowe TM. tRNAscan-SE 2.0: Improved detection and functional classification of transfer RNA genes. Nucleic Acids Research. 2021;49.

[56] Sella G, Hirsh AE. The application of statistical physics to evolutionary biology. Proceedings of the National Academy of Sciences of the United States of America. 2005 7;102:9541–6. Available from: https://www.pnas.org/content/102/27/9541https://www.pnas.org/content/102/27/9541.abstract.

[57] Nenadic O, Greenacre M. Correspondence Analysis in R, with two- and three-dimensional graphics: The ca package. Journal of Statistical Software. 2007;20:1–13.

[58] Maechler M, Rousseeuw P, Struyf A, Hubert M, Hornik K. cluster: Cluster Analysis Basics and Extensions; 2019.

[59] Thompson DA, Roy S, Chan M, Styczynsky MP, Pfiffner J, French C, et al. Evolutionary principles of modular gene regulation in yeasts. eLife. 2013 6;2:e00603. Available from: https://elifesciences.org/articles/00603.

[60] Revell LJ. Phylogenetic signal and linear regression on species data. Methods in Ecology and Evolution. 2010 12;1:319–29. Available from: http://doi.wiley.com/10.1111/j.2041-210X.2010.00044.x.

