## Supplementary material for "Macroevolutionary changes in natural selection on codon usage reflects evolution of the tRNA pool across a budding yeast subphylum": SI Appendix

### Supplemental Materials and Methods

#### Estimating unnormalized weights for the tRNA adaptation index

The unnormalized weights $W$ for the tRNA adaptation index (tAI) were estimated using the standard formula

$$W_i = \sum_{j=1}^{n_i} (1 - \mathrm{WP}_{j:i})\mathrm{tGCN}_{ij} \tag{1}$$

where the weight for codon $i$ is a function of the number of tGCN $n_i$ that recognize $i$ through Watson-Crick or wobble base pairing. In the case of wobble base pairing, a wobble penalty $\mathrm{WP}_{j:i}$ is used to reflect a decrease in the efficiency anticodon-codon base pairing. We used wobble penalites estimated previously for *S. cerevisiae*: $\mathrm{WP}_{G:U} = 0.41$, $\mathrm{WP}_{I:C} = 0.28$, $\mathrm{WP}_{I:A} = 0.9999$ , and $\mathrm{WP}_{U:G} = 0.68$. Note that $I$ indicates the inosine modification (A to I) at the $34^{th}$ nucleotide of a tRNA. We note that in the case of the CTG-Ser1 and CTG-Ser2 species, we treated the tGCN for CTG codons as 0.

#### Comparing selection coefficients $\Delta\eta$ with tRNA modification gene expression

The genes encoding tRNA modifications enzymes that make up the catalytic center of the Elongator Complex – IKI3 (ELP1), ELP2, and ELP3 – were identified in each Saccharomycotina yeasts tRNA modifications known to impact translation efficiency using a previously determined list of orthogroups (1). The relevant ROC-SEMPPR gene expression estimates $\phi$ were obtained for each gene and were compared to the selection coefficients $\Delta\eta$ for AAA, GAA, and CAA using multivariate phylogenetic regressions assuming a Pagel's $\lambda$ model as implemented in the R package **phylolm**. In addition to the effects of each of the Elongator Complex proteins, we also included the effects of the corresponding tGCN (specifically, log(tGCN)) for each of the codons and the genome-wide GC% in our regressions, i.e. $\Delta\eta_{Codon} \sim \phi_{IKI3} + \phi_{ELP2} + \phi_{ELP3} + tGCN_{Codon} + GC\% + s_{\phi} + Intercept$. Each independent variable was transformed into a Z-score to make the effects of each variable more comparable. A similar analysis was performed using the TAD2 and TAD3 proteins, which are responsible for the inosine modification.

### Supplemental Text

#### Estimates of stabilizing selection are stronger for mutation biases $\Delta M$ compared to selection coefficients $\Delta\eta$, on average

It is often assumed that a weaker phylogenetic signal is due to higher evolutionary rates. As stabilizing selection toward a single optimum value degrades phylogenetic signal (2), we fit an Ornstein-Uhlenbeck (3) model of trait evolution via the R package **geiger** (4) to the codon-specific parameters (using the standard deviation of the posterior distribution as measurement error), with the strength of stabilizing selection $\alpha$ compared between selection and mutation bias estimates using a Wilcox rank sum test. For the VarMut species, we performed these analyses using mutation bias estimates from either the Lower GC3% set or the Higher GC3% set.

We found the strength of stabilizing selection $\alpha$ is generally greater for estimates of mutation biases than estimates of selection (median $\alpha_{\Delta M_{\text{Lower GC3\%}}} = 0.011$, $\alpha_{\Delta\eta} = 0.005$, Wilcox rank sum test $p = 3.051E - 9$). We note the highest $\alpha$ values were for estimates of natural selection (Figure S10A). We obtain a similar result if using mutation bias estimates from the Higher GC3% set (Figure S10B). This is consistent with stronger stabilizing selection acting on the factors shaping mutation biases relative to natural selection, resulting in a stronger degradation of phylogenetic signal.

#### Potential super wobbling of tRNA translaing proline in the Saccharomycotina subphylum

Surprisingly, we found numerous codons do not have a corresponding tRNA based on the identified tRNA genes and standard wobble rules, implying inefficient elongation of these codons. Although many of these were likely spuriously missed tRNA genes by tRNAScan-SE, we note a lack of tRNA genes recognizing proline codons CCC/CCT in the clades CUG-Ser2 (2/4 species), Phaffomycetaceae (34/34 species), Pichiaceae (46/61 species), and Saccharomycodaceae (8/8) species (Figure S14). A gene encoding Pro-tRNA$^{\mathrm{UGG}}$codon is present in each of these species, but not at appreciably different amounts than observed in the other species (Welch two-sample t-test, $p = 0.7742$). Assuming the corresponding tRNA genes are truly missing in these clades, this suggests the occurrence of super-wobbling for proline, whereby an unmodified U34 allows a tRNA to bind all 4 codons (5).

#### The average strength of selection on codon usage significantly correlates with total number of tRNA genes

Selection for fast growth is hypothesized to shape the overall strength of translational selection, particularly in microbes (6; 7; 8; 9). For example, organisms with more translational resources (e.g., tRNA genes) are hypothesized to be under stronger selection for fast growth, resulting in stronger selection for efficient translation. We found the mean absolute selection coefficients per species, which reflects the average strength of selection across all codons, was positively correlated with the total number of tRNA genes per genome (Spearman rank correlation $\rho = 0.22$, $p = 0.00025$, Figure S15A). This conflicts with (10), who found no significant relationship between $S$ (11) (based on tAI and the effective number of codons, not to be confused with the population genetics parameter $S$) and the total number of tRNA genes after accounting for shared ancestry. We note $S$ is not a theoretically justified estimate of natural selection on codon usage (11; 7); perhaps unsurprisingly, mean absolute selection coefficients were only weakly correlated with $S$ (Spearman rank correlation $\rho = 0.13$, $p = 0.035$, Figure S15B). Our results suggest natural selection on codon usage increases slightly with investment in translational resources, but other factors likely contributed to differences in adaptive CUB.

#### Mutation bias variation reflect changes to genome-wide GC% content

Genome-wide GC% varies across species, but it is unclear to what extent this variation is due to variation in mutation biases. For example, mutations appear universally AT-biased in bacteria with variation in GC% due to natural selection or biased gene conversion (12). In contrast to bacteria, more recent work found no universal mutation bias in the Saccharomycotina subphylum (13). We compared our estimates of mutation bias $\Delta M$ to genome-wide GC% across species. We focused on the ConstMut and the Lower GC3% estimates, as these better predict genome-wide GC% (Figure S8). To assess the impact of evolutionary processes that may favor GC-ending codons over AT-ending codons, mutation biases were modified such that all GC-ending codons were relative to the corresponding AT-ending codon. This is the similar to the "transition selection coefficients" presented in the main text. As ROC-SEMPPR's selection coefficients for each amino acid are estimated relative to a pre-defined reference (the alphabetically last codon), we calculated the "transition mutation biases"as $\Delta M_{\mathrm{Transition}}$ as

$$\Delta M^{R-R}_{\mathrm{Transition}} = \Delta M_{\mathrm{NNG}} - \Delta M_{\mathrm{NNA}} \quad (2)$$

$$\Delta M^{Y-Y}_{\mathrm{Transition}} = \Delta M_{\mathrm{NNC}} - \Delta M_{\mathrm{NNT}} \quad (3)$$

to estimate the relative strength and direction of mutation bias between purines (NNA vs. NNG) and pyrimidines (NNC vs. NNT). In this context, a positive mutation bias indicates the GC-ending codon is disfavored by mutation bias relative to the AT-ending codon. Similarly, a negative value indicates the GC-ending codon is favored relative to the AT-ending codon. Our mutation biases were well-correlated with GC% across species (Figure S22). The correlation between mutation biases and genome-wide GC% is consistent with across-species variation in GC% being shaped by differences in mutation bias.

##### Across-species variation in GC% weakly reflects variation in mismatch-repair genes

Mutation biases $\Delta M$ are correlated with GC% aacross species, but what molecular factors drive shifts in mutation biases? Previous work found the loss of mismatch repair (MMR) genes were correlated with changes to GC% across fungi, specifically genomes that have experienced fewer losses of MMR genes had higher GC%, on average (14). Using a previously published presence-absence MMR gene matrix (14), species with the greatest number of missing mismatch repair genes have a lower genome-wide GC%; however, this constituted only 8 of the 293 species (Figure S23A). Overall, we find no statistically significant relationship between the number of missing MMR genes and genome-wide GC% after controlling for phylogeny (PGLS slope $\beta = -0.0023$, $p = 0.25$). Although species with a relatively large number of missing MMR genes have a stronger AT-bias, the variation observed in GC% across species is poorly explained by the number of MMR genes.

As an alternative, we examined if changes to gene expression $\phi$ of MMR genes are correlated with changes to GC%. Single-copy orthologous MMR genes were identified using orthogroups forom (1). Overall, very few MMR protein-coding sequences show statistically significant correlations between gene expression and GC%, with the exceptions of MSH3, MSH6, POL3, and RFC2 (Figure S23B). Except for MSH3, these genes' expression levels are negatively correlated with GC%, indicating that as expression increases, these species become more AT-biased. Based on the assumption that fewer MMR genes leads to a stronger AT-bias in fungi, as proposed in (14), these negative correlations are counterintuitive; however, these correlations are weak. The overall lack of relationship between GC% and gene expression of MMR proteins could be due to interactions between changes in gene expression of different MMR genes. We performed a phylogenetic principal component analysis (phylo-PCA) based on the predicted gene expression values of the MMR genes using the R package **phytools**. The first principal component axis is negatively correlated with GC% across species, but this correlation is weak (Figure S23C, Spearman rank correlation $\rho = -0.16, p = 0.0047$). Based on the loadings, the phylogenetic-PCA suggests a different subset of genes may be contributing to across-species variation in GC% compared to the individual gene analysis, particularly CDC17 and RFC5 (Figure S23D). Both analyses suggest a role for shifts in the expression of MMR genes in shaping mutation biases, but the exact relationship between these variables remains unclear.

#### Supplemental Figures

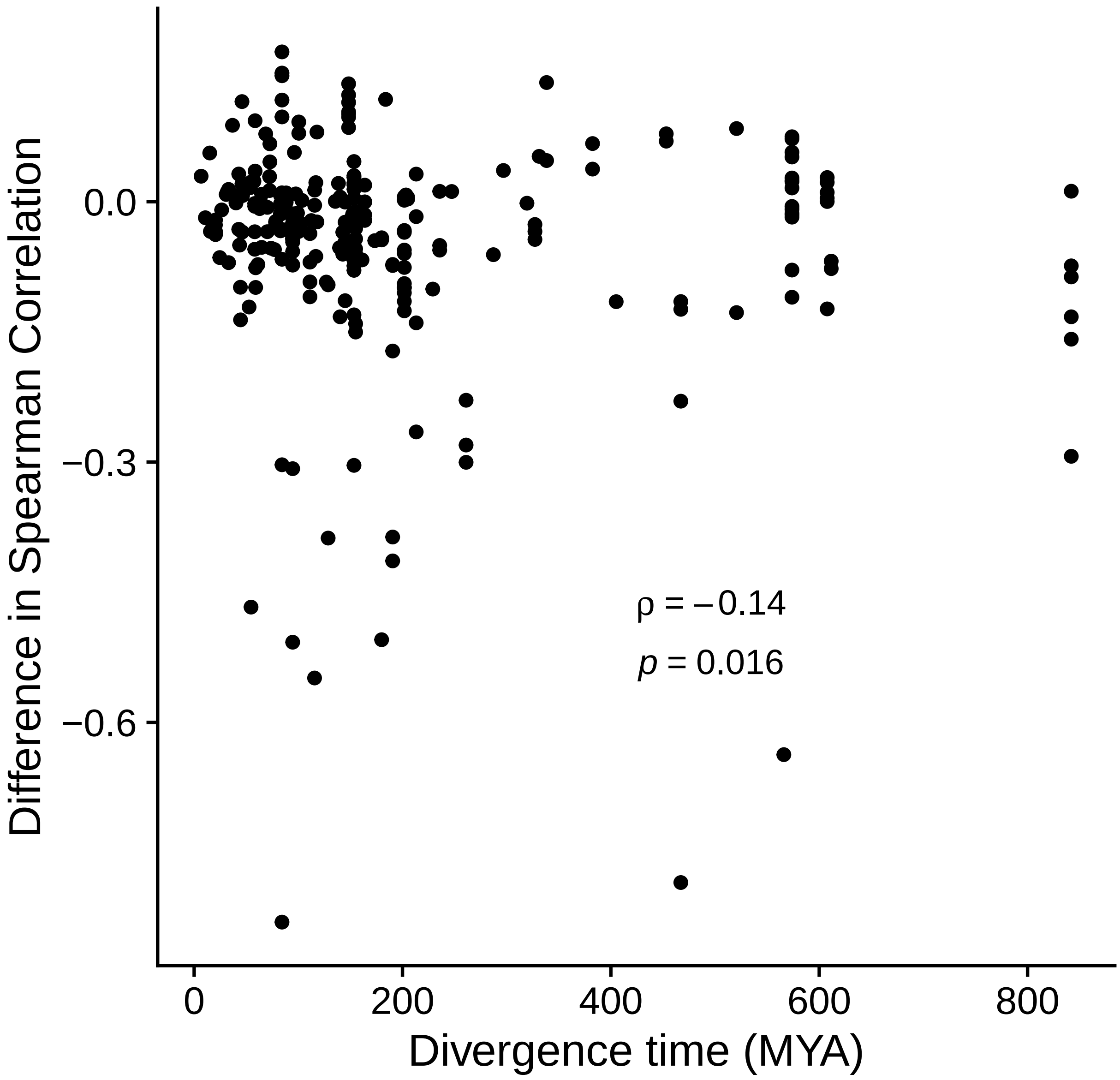


Figure S1: Comparison of Spearman correlation between ROC-SEMPPR $\phi$ and empirical gene expression for the target species and the reference species (i.e. the species for which the empirical measurements were taken).

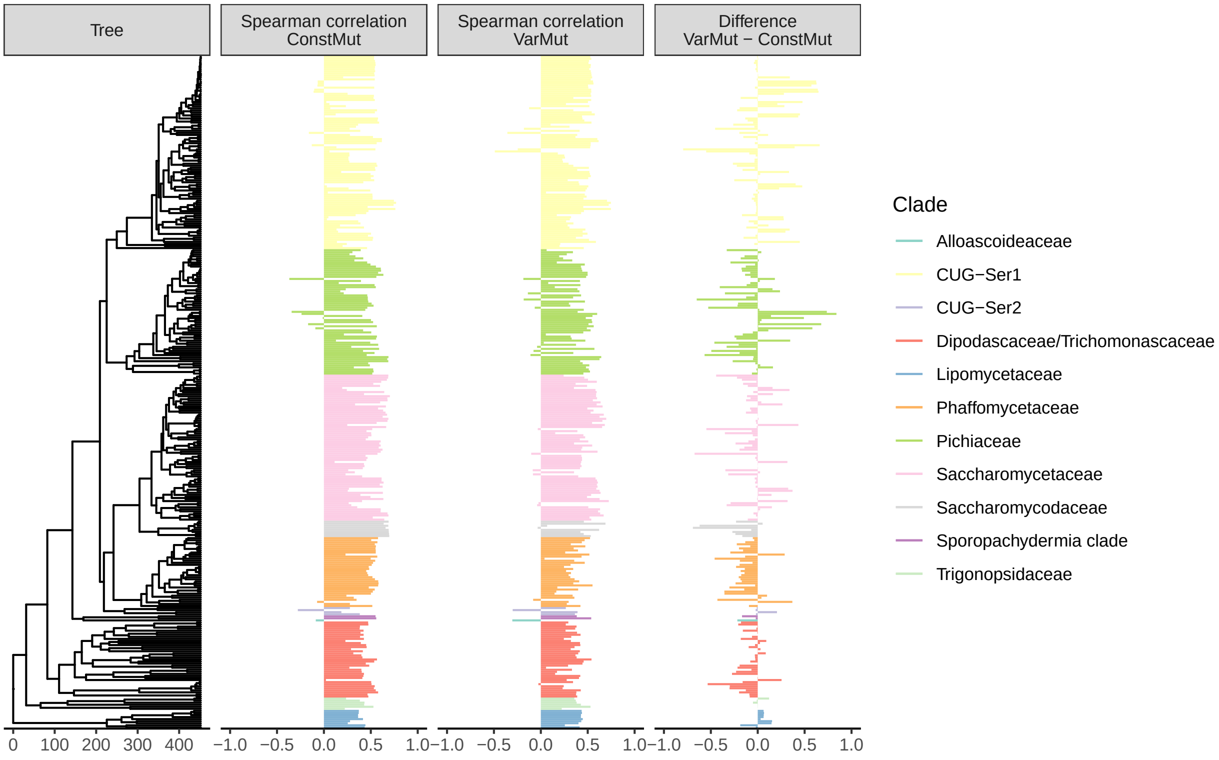


Figure S2: Spearman rank correlation $\rho$ between ROC-SEMPPR predicted gene expression $\phi$ and empirical gene expression (RNA-seq) for the ConstMut and VarMut model fits across the 327 budding yeasts.

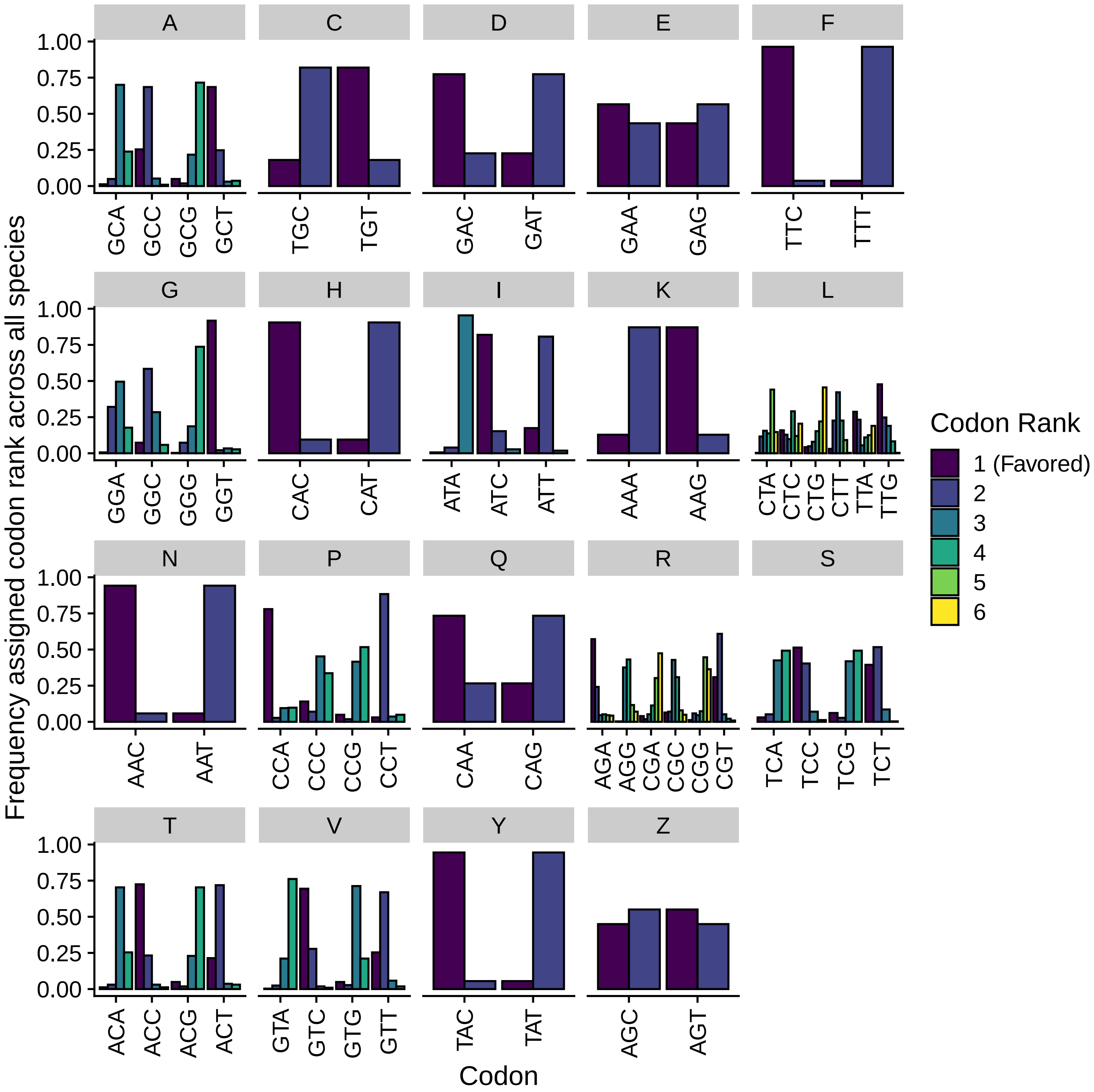


Figure S3: Distribution of codon rankings across 327 budding yeasts based on selection coefficients $\Delta\eta$, with a rank of 1 indicating a codon is the "best" codon, i.e. the one favored by natural selection for a given set of synonymous codons. "Z" indicates the serine codons AGC/AGT, which are treated as separate from the other 4 serine ("S") codons.

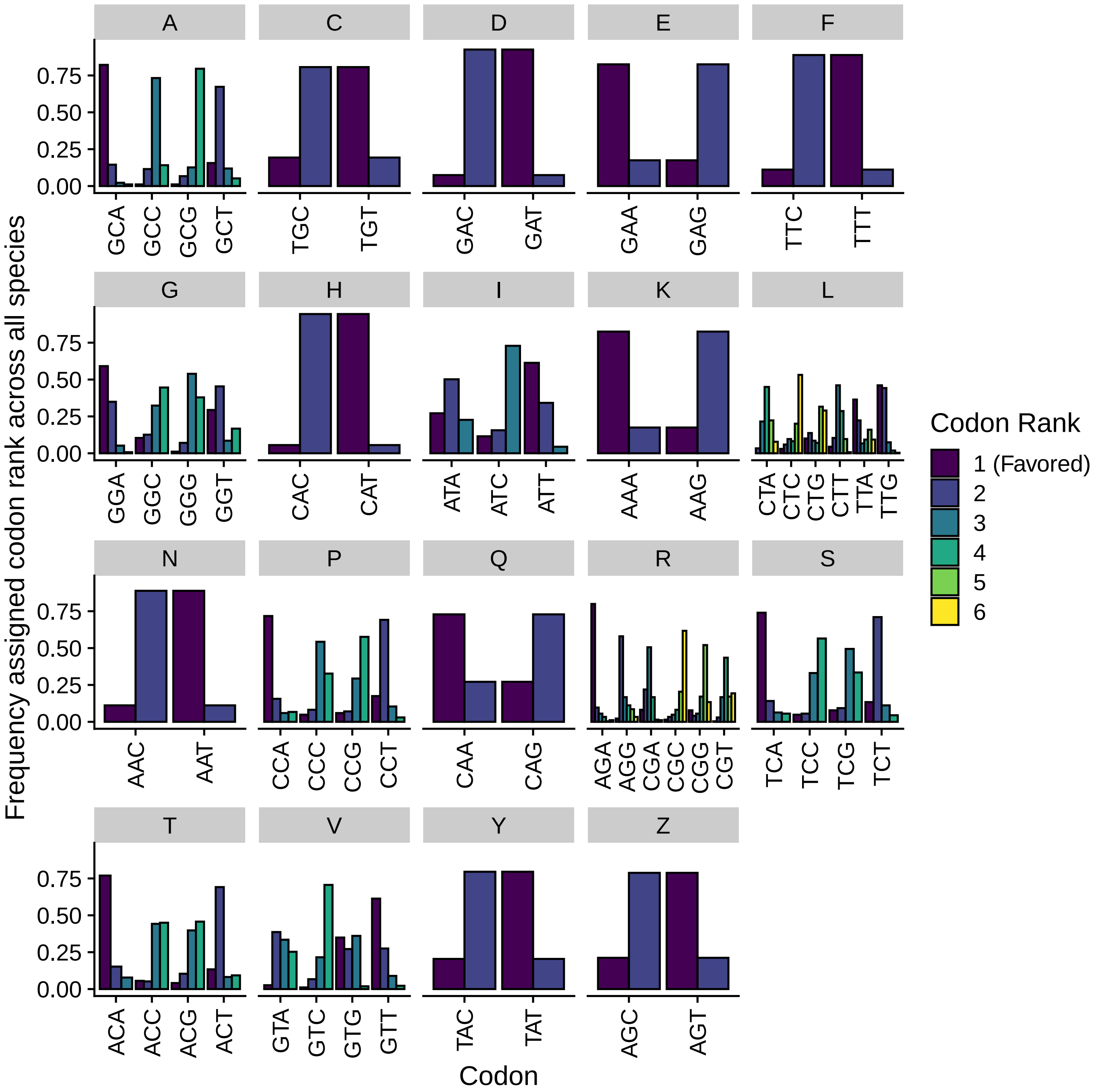


Figure S4: Distribution of codon rankings across 327 budding yeasts based on mutation biases $\Delta M$ for the ConstMut species, with a rank of 1 indicating a codon is the "best" codon, i.e. the one favored by mutation bias for a given set of synonymous codons. "Z" indicates the serine codons AGC/AGT, which are treated as separate from the other 4 serine ("S") codons.

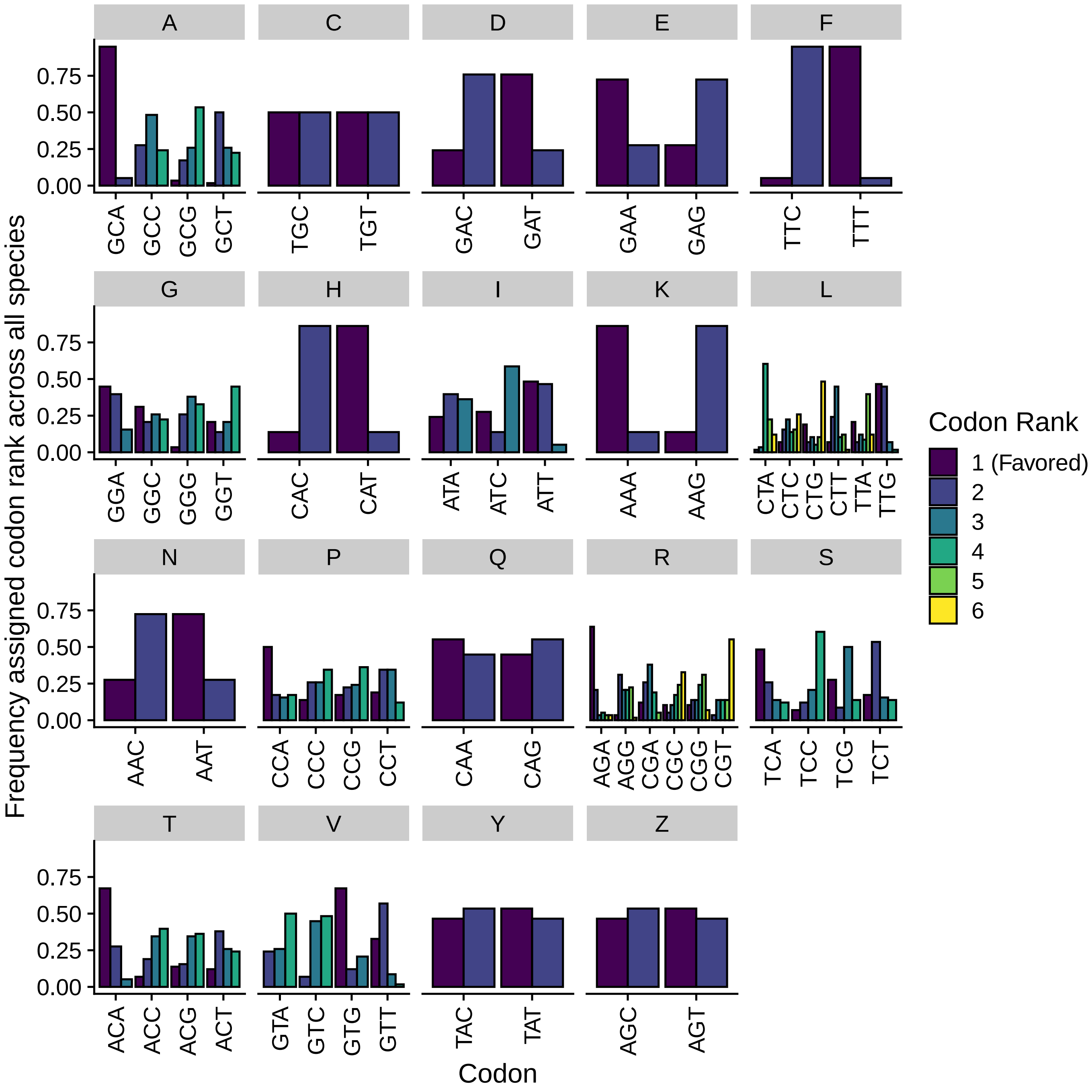


Figure S5: Distribution of codon rankings across 327 budding yeasts based on mutation biases $\Delta M$ for the Lower GC3% set of the VarMut species, with a rank of 1 indicating a codon is the "best" codon, i.e. the one favored by mutation bias for a given set of synonymous codons. "Z" indicates the serine codons AGC/AGT, which are treated as separate from the other 4 serine ("S") codons.

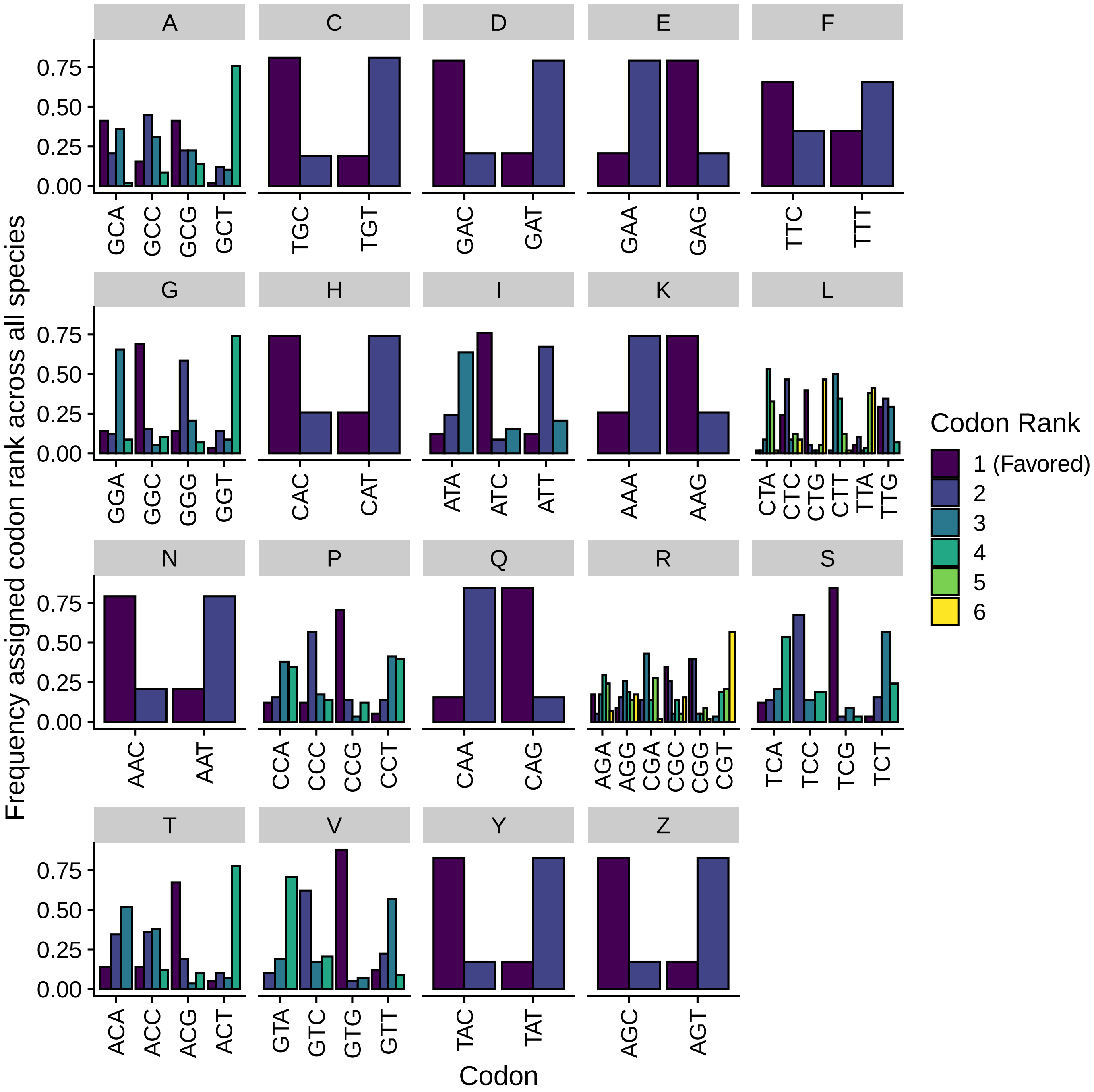


Figure S6: Distribution of codon rankings across 327 budding yeasts based on mutation biases $\Delta M$ for the Higher GC3% set of the VarMut species, with a rank of 1 indicating a codon is the "best" codon, i.e. the one favored by mutation bias for a given set of synonymous codons. "Z" indicates the serine codons AGC/AGT, which are treated as separate from the other 4 serine ("S") codons.

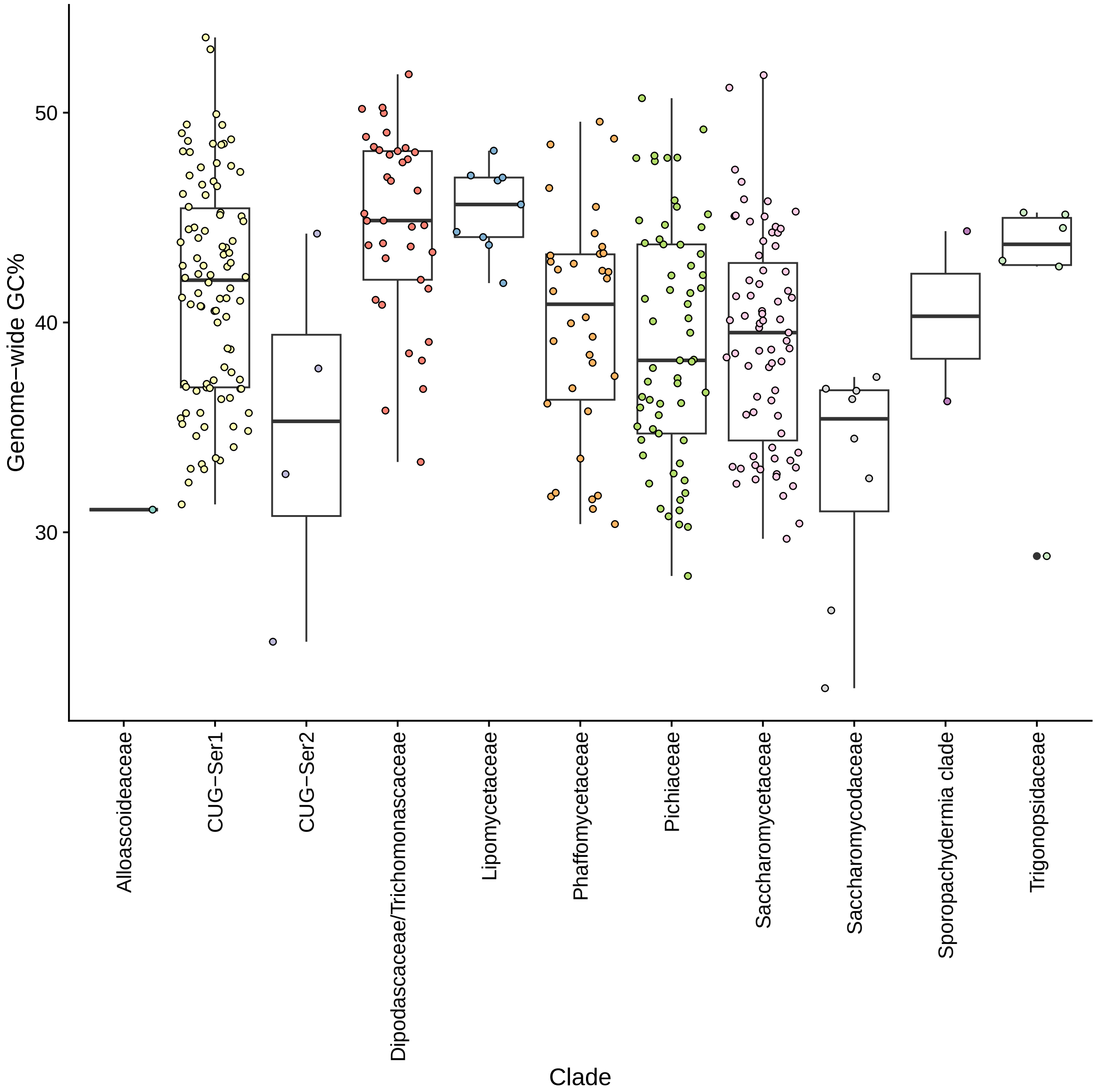


Figure S7: Boxplots showing distribution of genome-wide GC% for all 327 species depending on major clade.

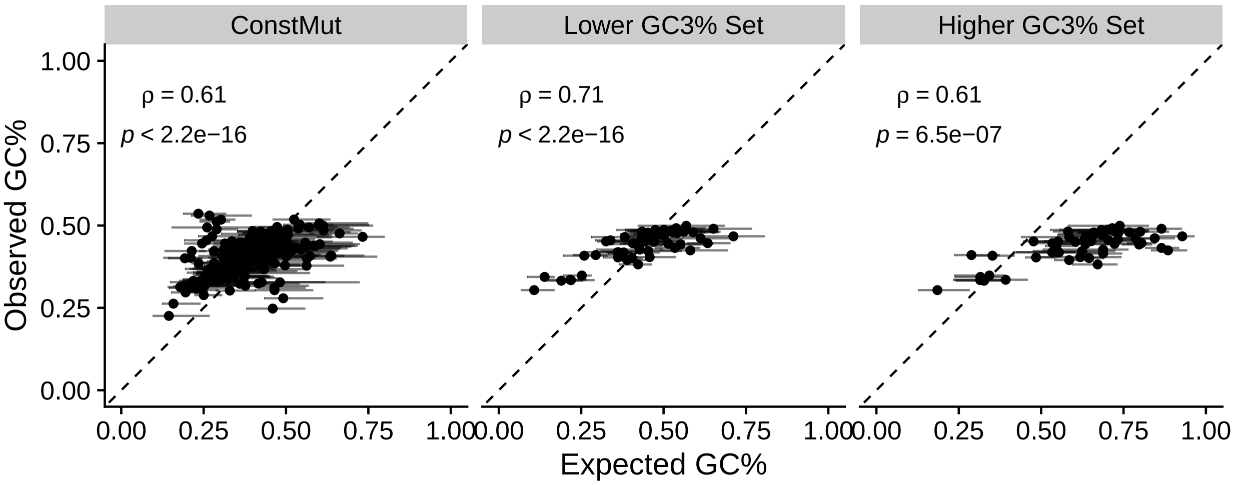


Figure S8: Comparison of median expected vs observed genome-wide GC% based on the mutation bias estimates of the 4-codon amino acids. Error bars represent the range of expected genome-wide GC% across the 4-codon amino acids. Spearman rank correlations $\rho$ and associated $p-$values are reported.

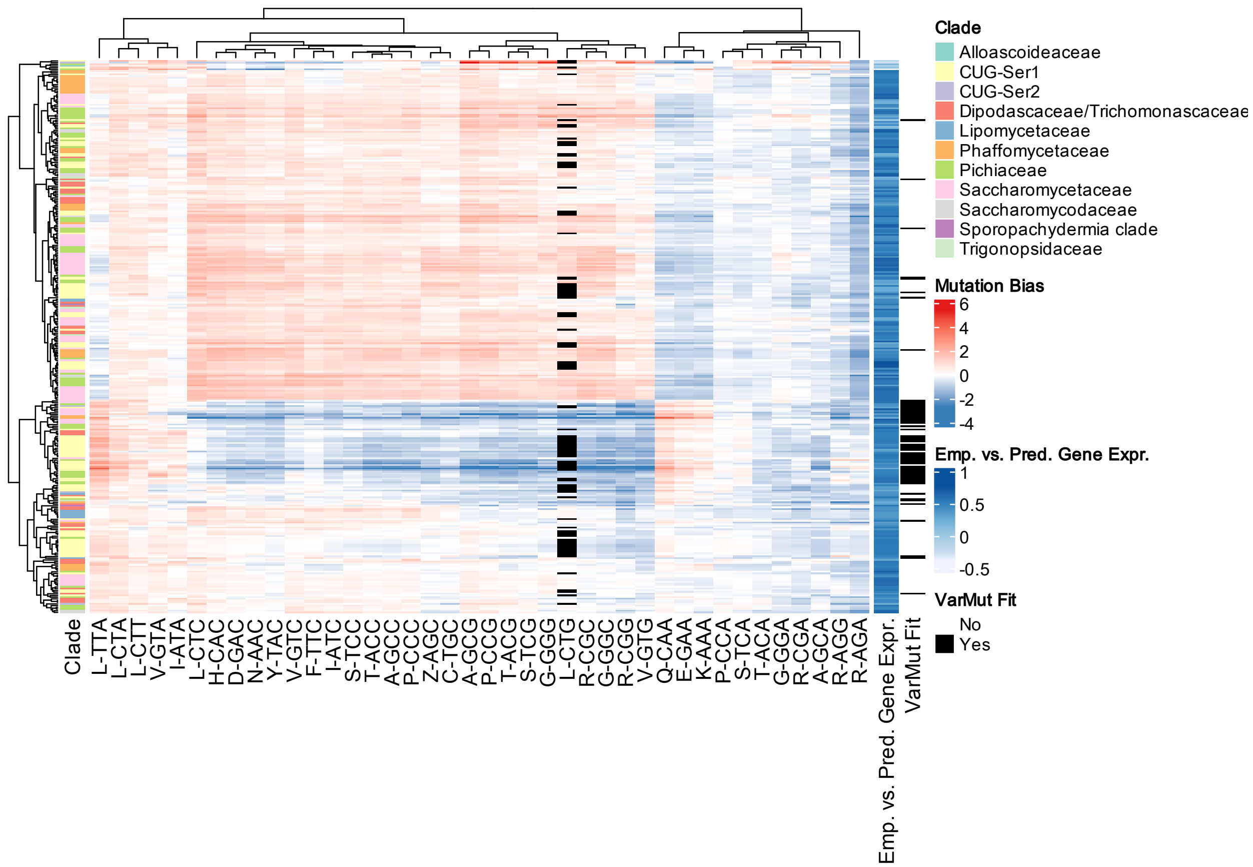


Figure S9: Hierarchical clustering of mutation biases $\Delta M$ across the 327 budding yeasts using estimates from the ConstMut and Higher GC3% set of the VarMut species. The "Clade" color bar indicates the major clade of the species. The "Emp.vs.Pred.Gene.Exp" color bar indicates the correlation of predicted gene expression $\phi$ with empirical RNA-seq data. "VarMut.fit" indicates if the species was better fit by the VarMut model. "Z" indicates the serine codons AGC/AGT, which are treated as separate from the other 4 serine codons.

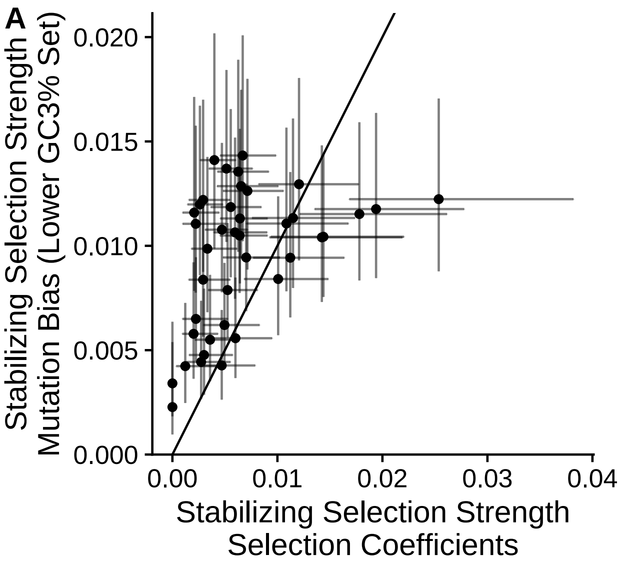


Figure S10: Comparison of stabilizing selection $\alpha$ parameters fit via an Ornstein-Uhlenbeck model to selection coefficients $\Delta\eta$ and mutation biases $\Delta M$ for each individual codon (reference codons excluded). Error bars indicate 95% confidence intervals. (A) Mutation biases from Lower GC3% set for species best fit by the VarMut model (ConstMut fit otherwise). (B) Mutation biases from Higher GC3% set for species best fit by the VarMut model (ConstMut fit otherwise).

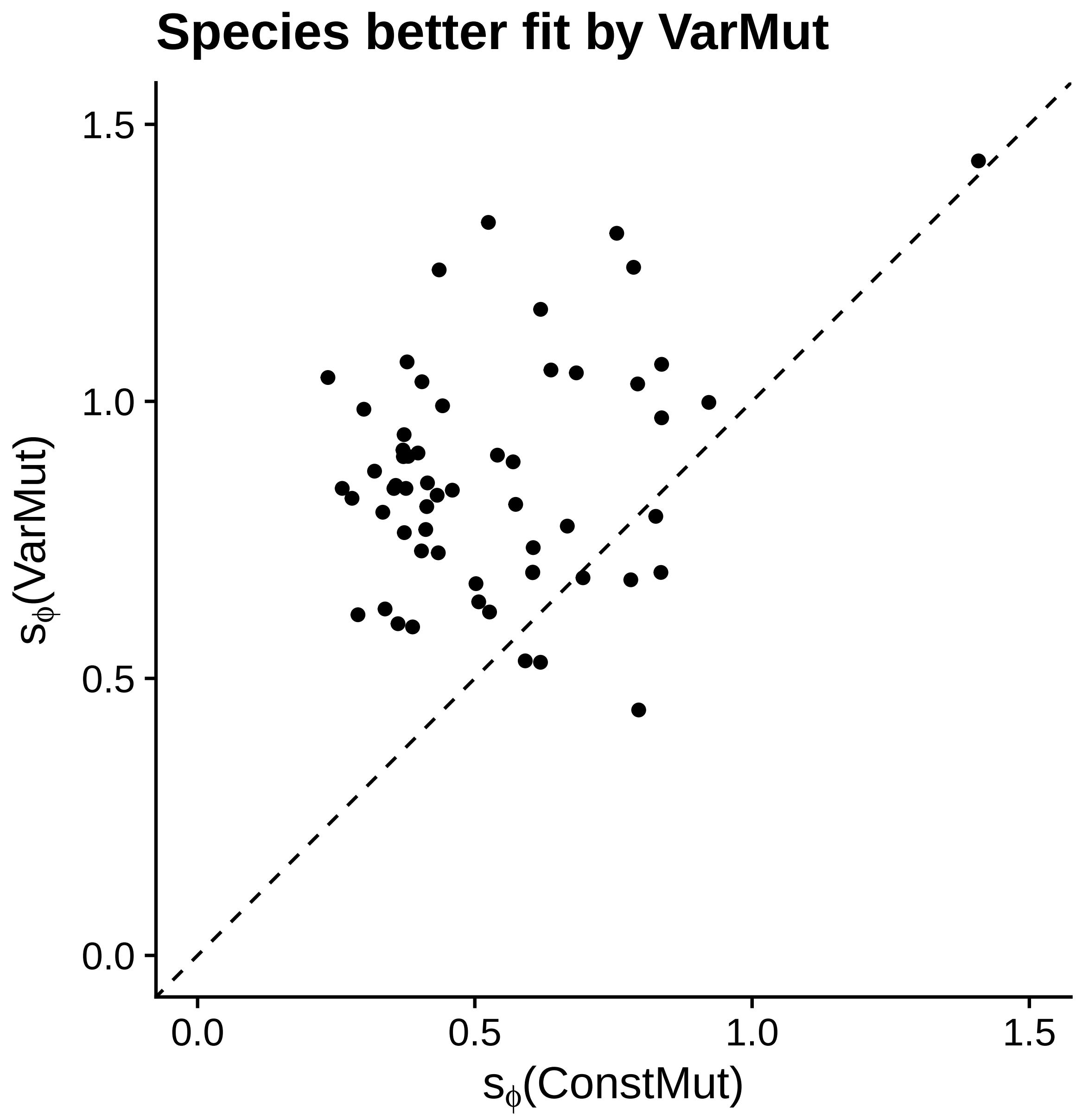


Figure S11: Comparison of $s_\phi$ estimates from the ConstMut and VarMut models for species better fit by the latter.

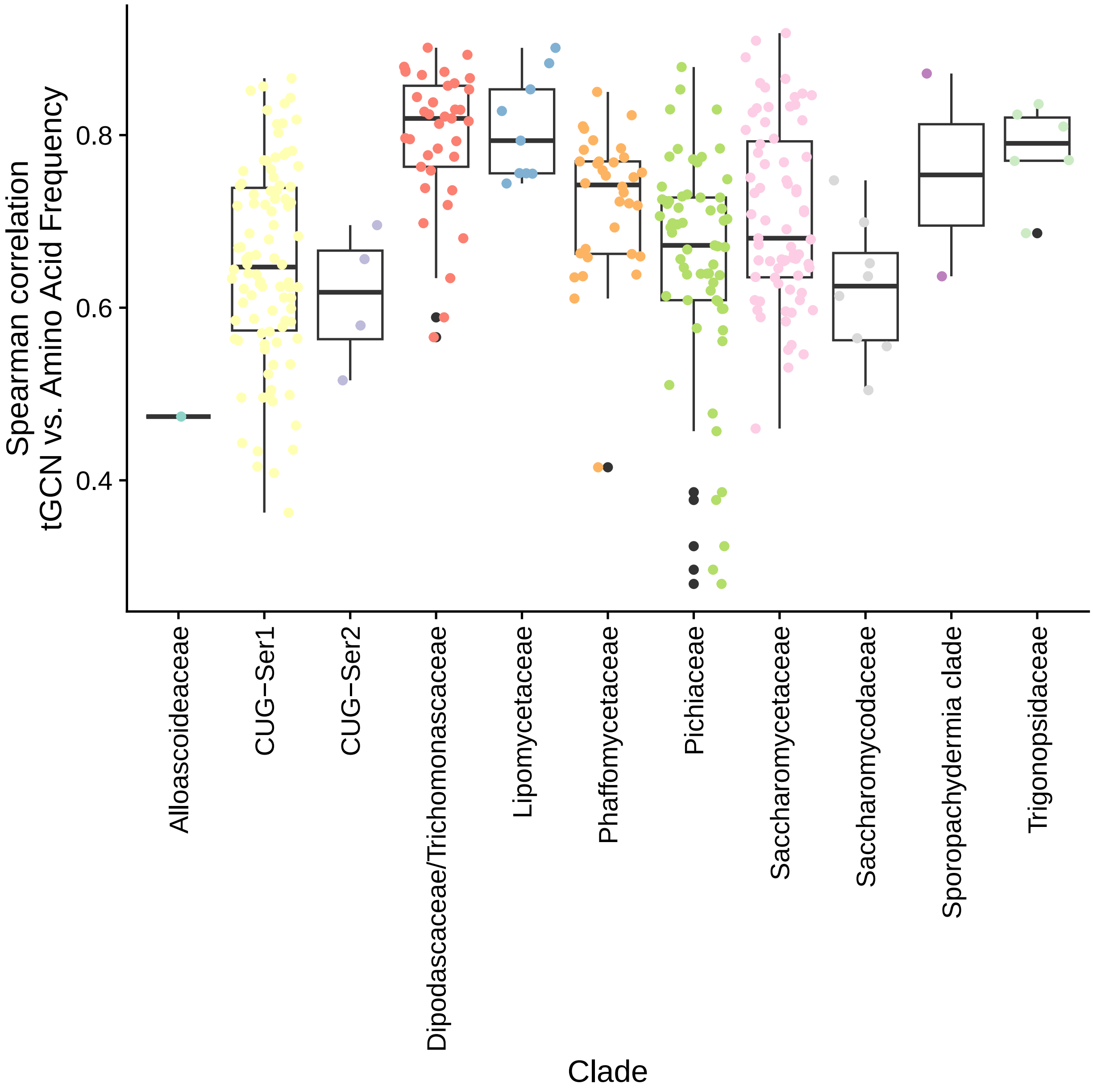


Figure S12: Distribution per clade of Spearman rank correlation $\rho$ between proteome-wide amino acid frequency and per-amino acid total tRNA gene copy number (tGCN).

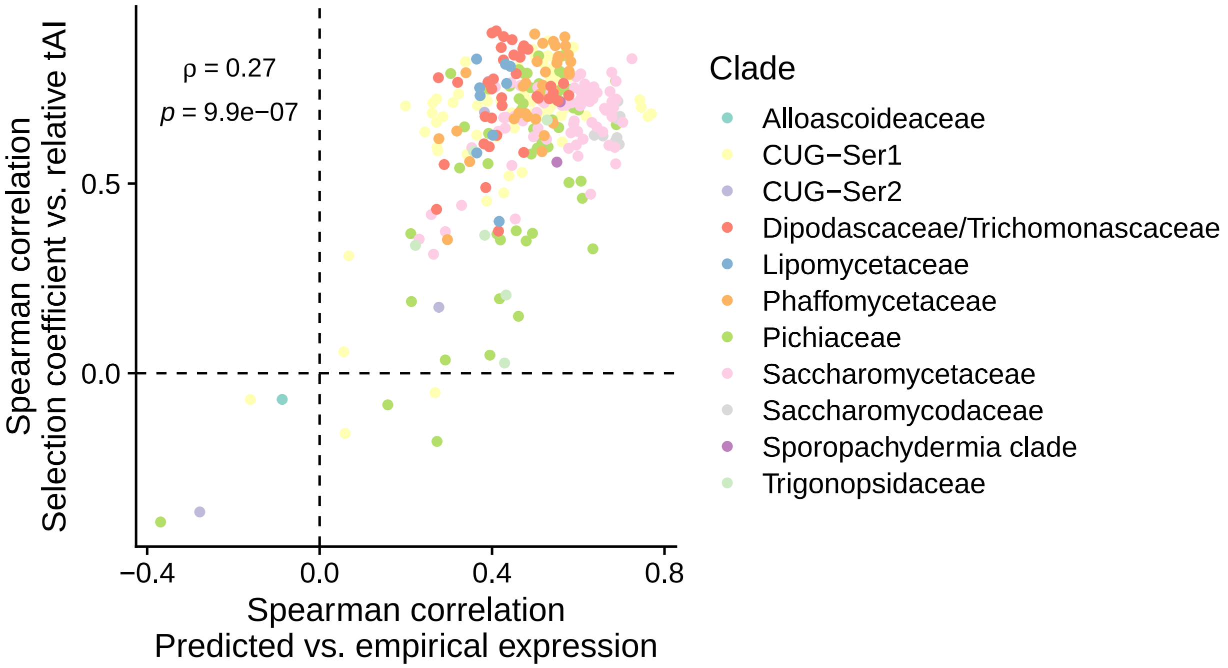


Figure S13: Scatter plot showing correlation the Spearman rank correlations $\rho$ of the empirical (RNA-seq TPM) vs. predicted gene expression (ROC-SEMPPR $\phi$) (x-axis) and the relative tAI $\Delta(1/\text{tAI})$ and selection coefficients $\Delta\eta$. Note that we did not apply phylogenetic indpendent contrasts or other approaches, as the goal here was to see how the correlations compared rather than to make an inference about evolution. Even so, the $p$-value should be taken with caution due to the non-independence between species.

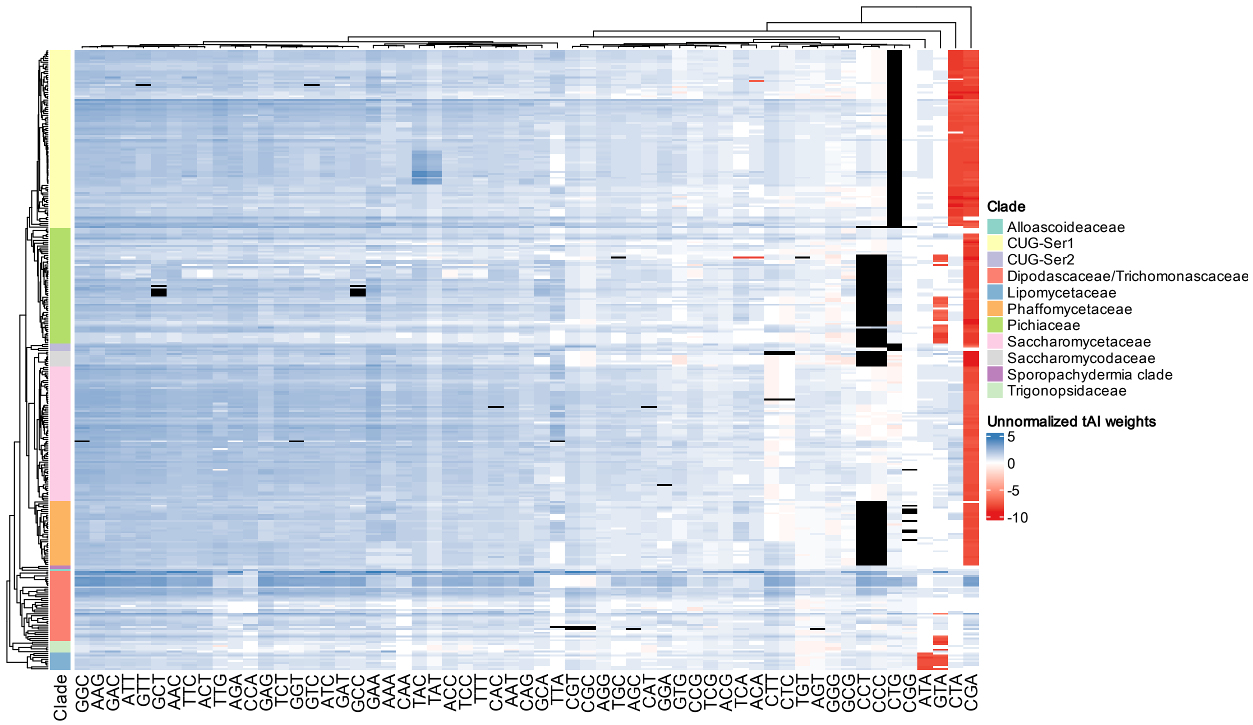


Figure S14: Heatmap of unnormalized weights used for tAI. The color ”black” indicates a weight could not be calculated for a codon based on the provided tRNA genes and standard wobble rules.

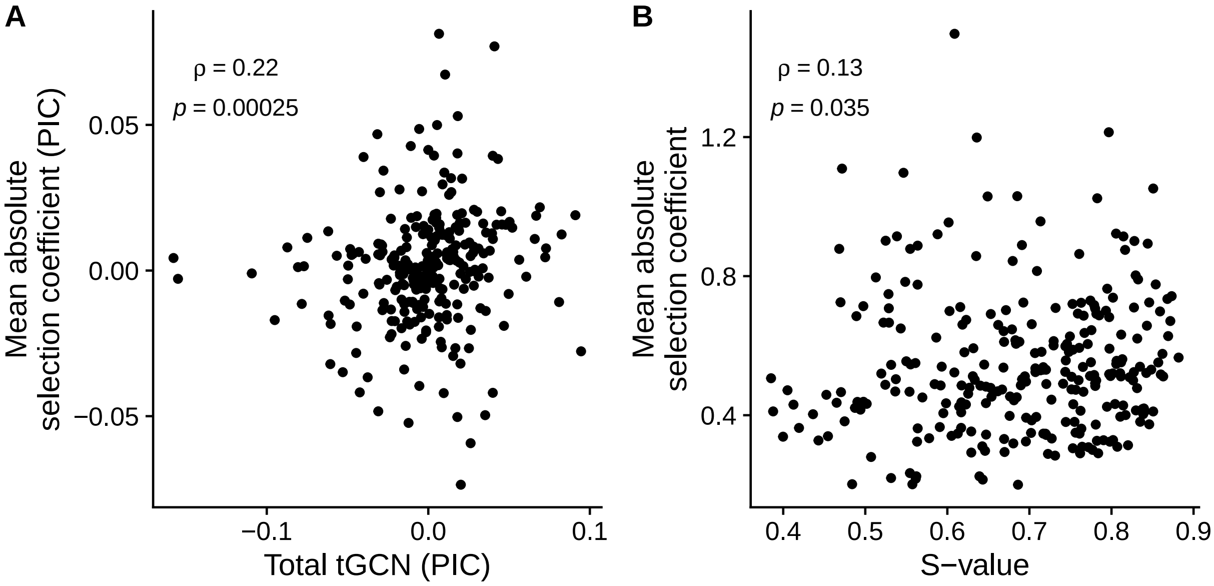


Figure S15: Comparison of mean absolute selection coefficient $|\Delta\eta|$ and the $S$ value. Spearman rank correlation $R$ is reported.

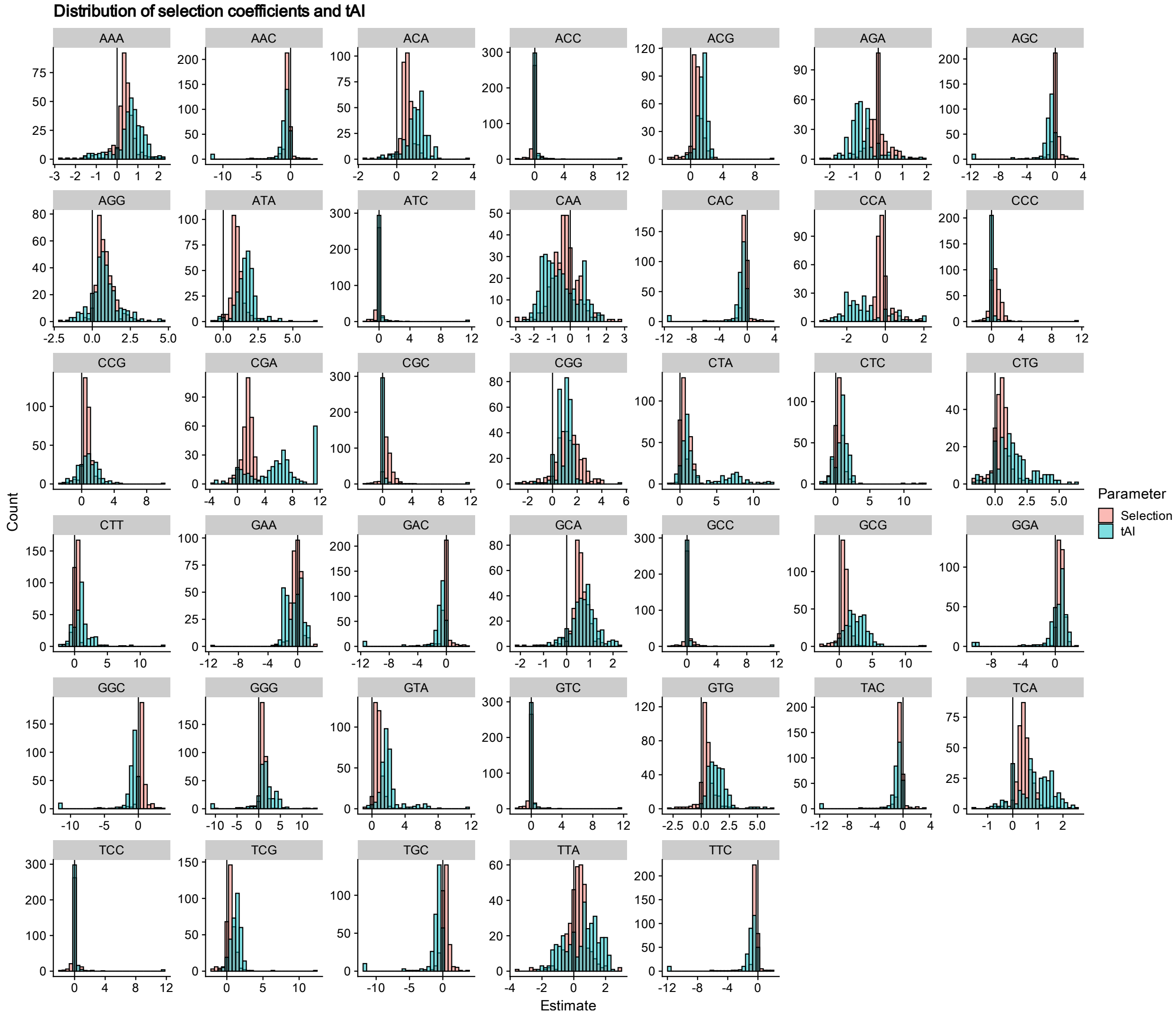


Figure S16: Distribution of selection coefficients $\Delta\eta$ and the relative tAI weights for all 327 Saccharomycotina yeasts.

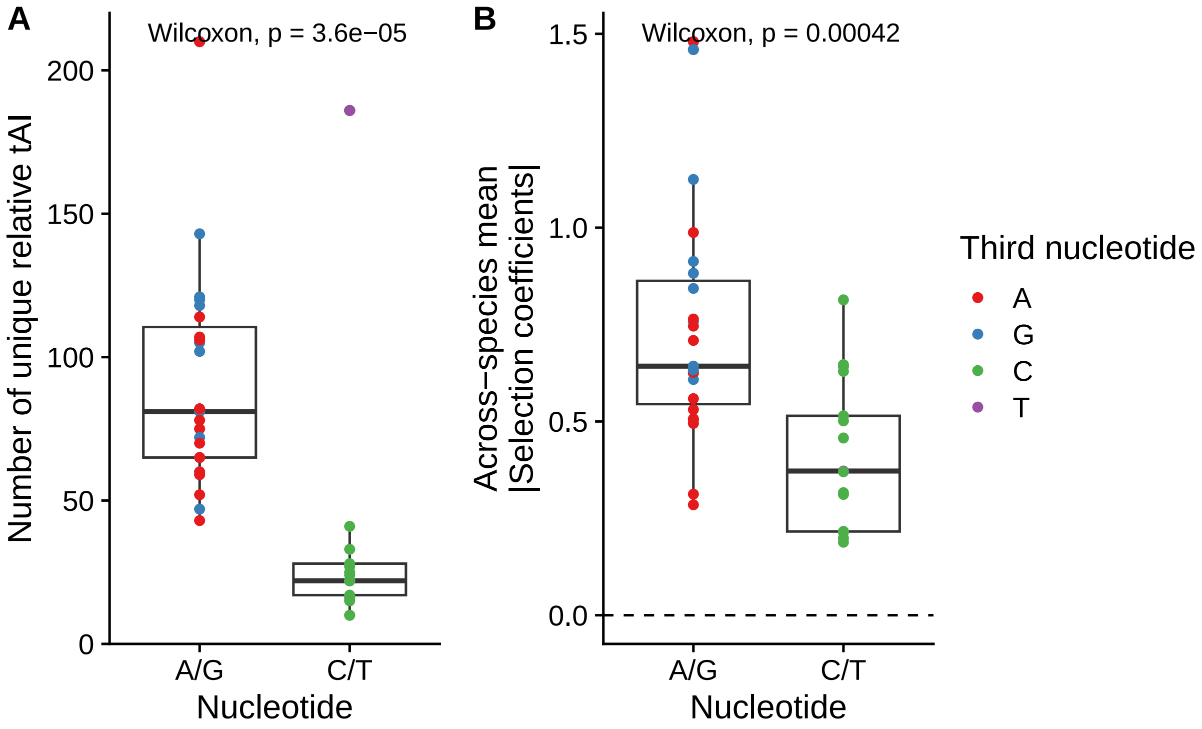


Figure S17: (A) Number of unique relative tAI values $\Delta 1/\text{tAI}$ across the Saccharomycotina subphylum after removing the suspected poorly fit species based on if the third nucleotide is A/G vs. C/T. The p-value of a Wilcox rank sum test is reported. (B) Across-species mean of the absolute selection coefficients $|\Delta\eta|$ based on if the third nucleotide is A/G vs. C/T. The p-value of a Wilcox rank sum test is reported.

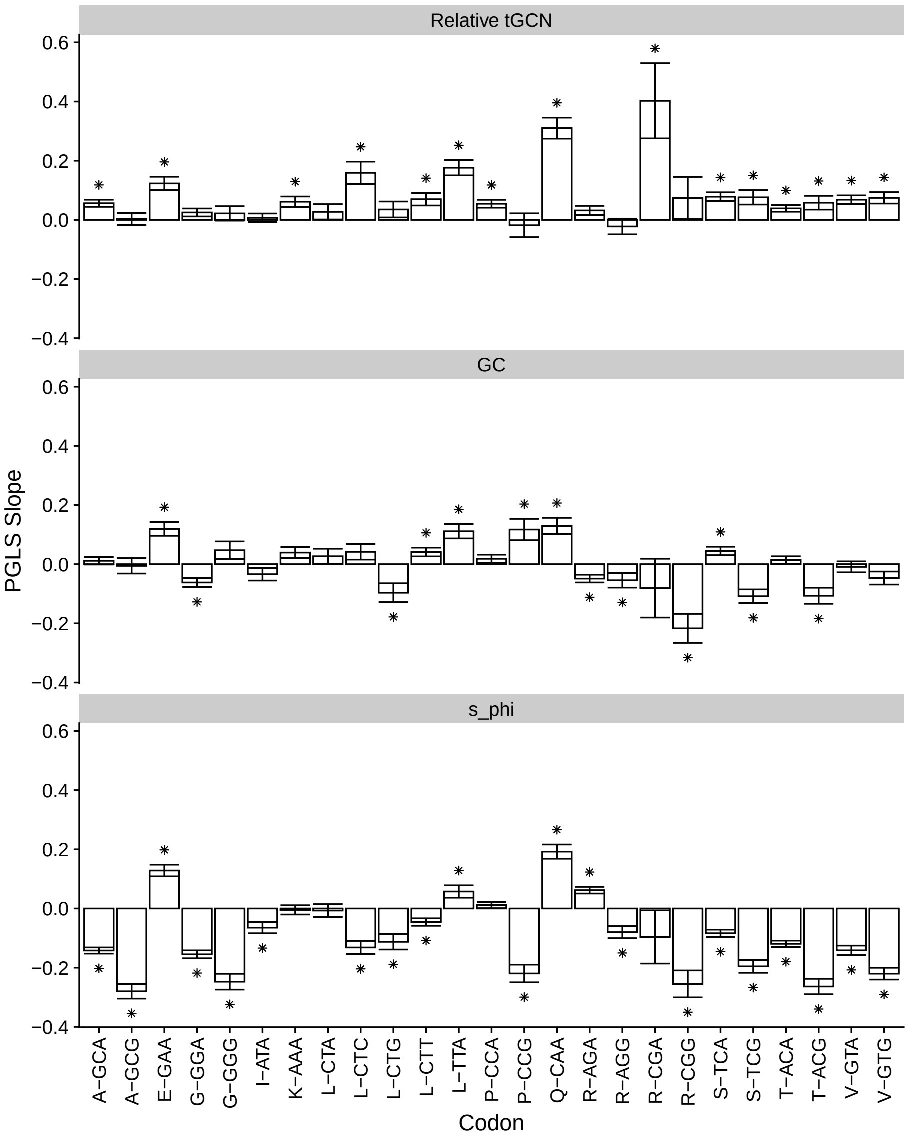


Figure S18: PGLS slopes (Pagel's $\lambda$ model of trait evolution) showing relationship of relative tAI $\Delta 1/\text{tAI}$, genome-wide GC%, genome size (as proxy for effective population size $N_e$), and $s_\phi$ (as a potential confounder) with selection coefficients $\Delta\eta$. Codons considered are those for which differences in elongation rates (using relative tAI as proxy) could be determined based on tGCN. Error bars indicated $\pm 1$ std. error. "*" indicates Benjamini-Hochberg adjusted $p < 0.05$.

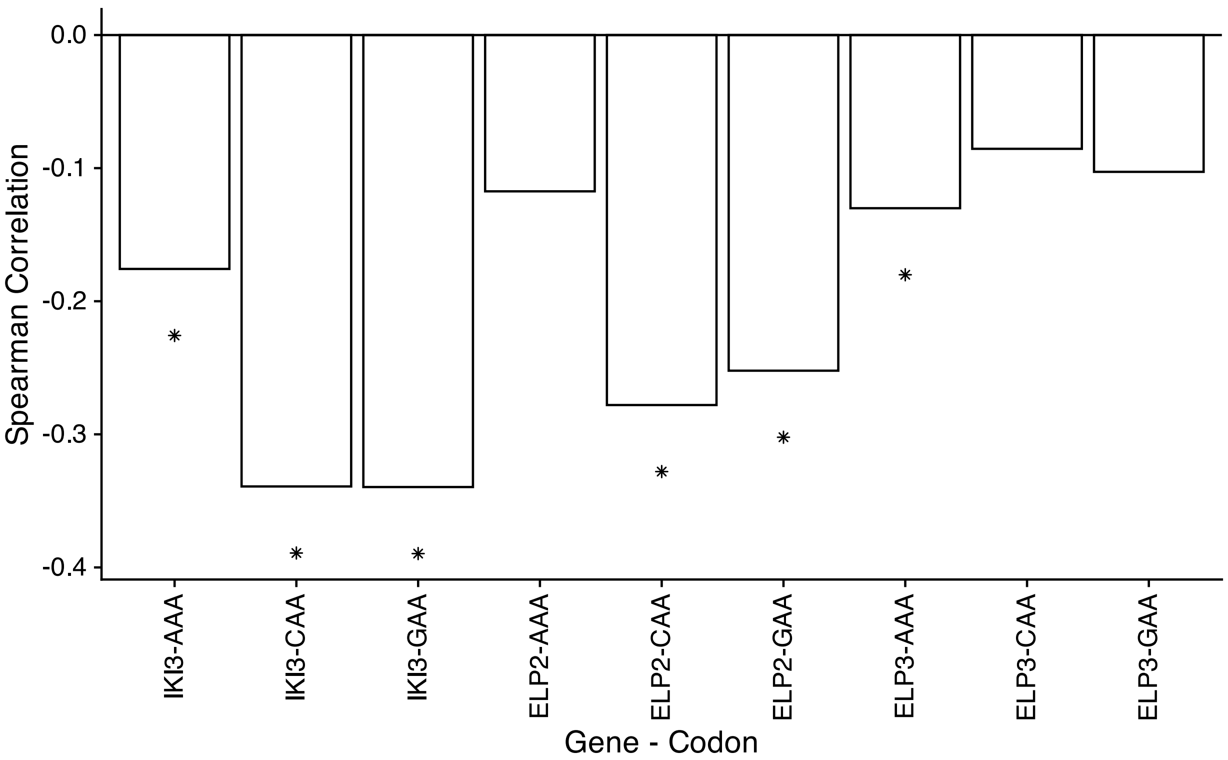


Figure S19: Barplots showing Spearman rank correlations between selection coefficients $\Delta\eta$ and predicted gene expression $\phi$ of Elongator Complex proteins IKI3, ELP2, and ELP3. Phylogenetic independent contrasts was applied before calculating correlations.

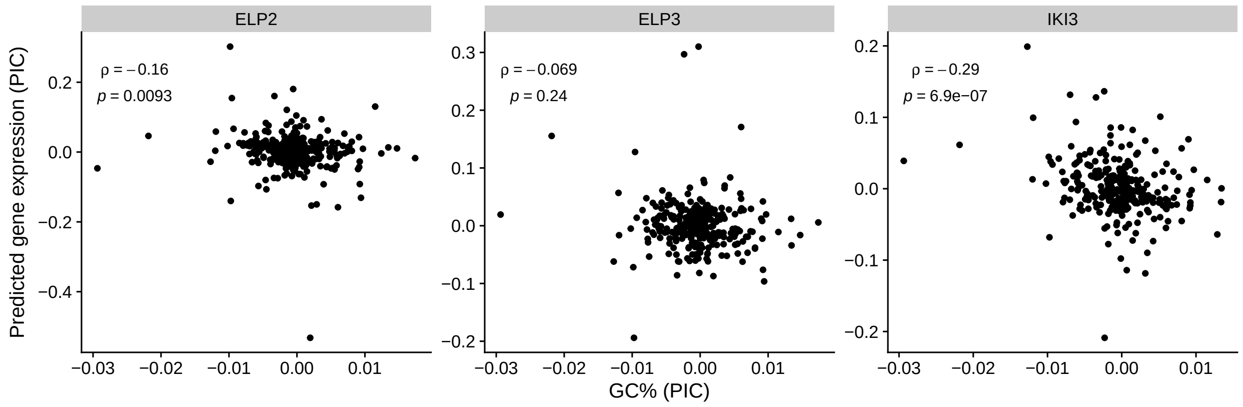


Figure S20: Scatter-plot showing the relationship between GC% and the predicted gene expression of the Elongator Complex proteins IKI3, ELP2, ELP3.

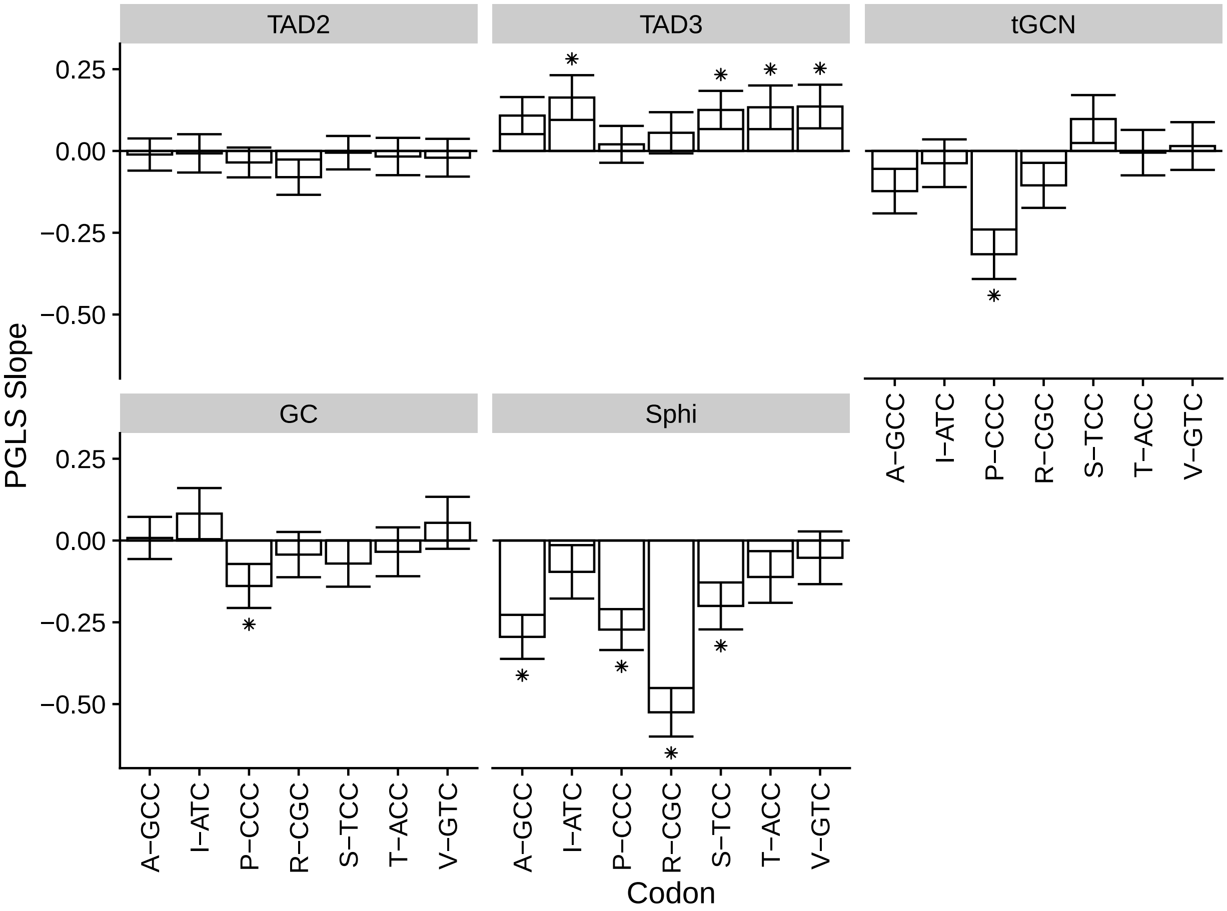


Figure S21: PGLS slopes (Pagel's $\lambda$ model of trait evolution) showing relationship of expression of tRNA modification enzymes involved in inosine modification, tGCN, genome-wide GC%, and $s_\phi$ (as a potential confounder) with selection coefficients $\Delta\eta$. Error bars indicated $\pm 1$ std. error. "*" indicates Benjamini-Hochberg adjusted $p < 0.05$.

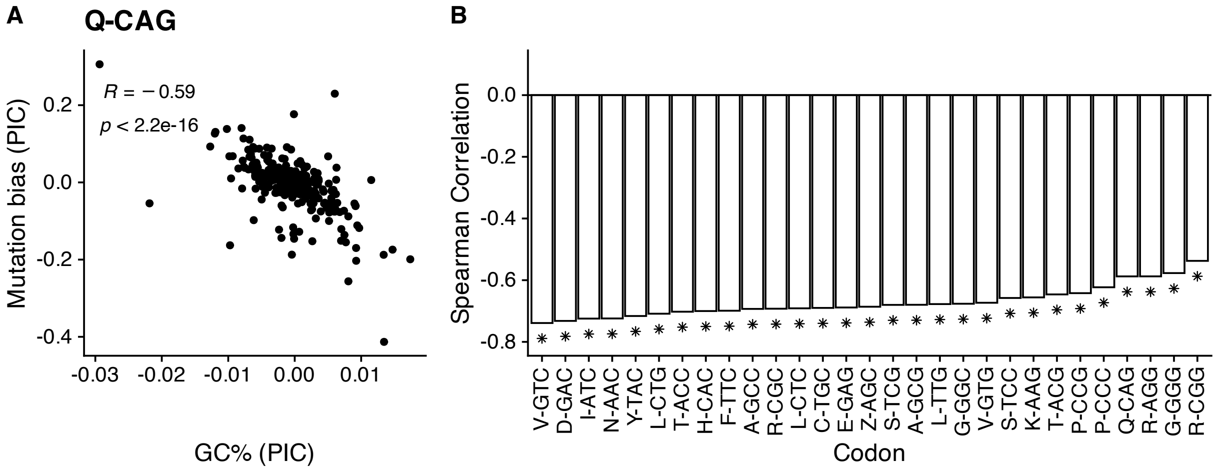


Figure S22: Relationship between GC% and mutation bias $\Delta M$ estimates across the phylogeny. For all cases, mutation biases were modified to reflect mutation biases between either A and G ending codons or C and T ending codons, as opposed to the alphabetically last codon (the default of AnaCoDa). Phylogenetic independent contrasts (PIC) were applied before calculating the Spearman rank correlations. Note that a negative value of mutation biases $\Delta M$ indicates a bias towards G/C-ending codons relative to A/T-ending codons (A) Example scatter-plot showing the relationship between GC% and mutation bias $\Delta M$ for codon CAG (relative to CAA). The negative correlation indicates that GC% increases as CAG becomes more favored by mutation relative to CAA. (B) Across-species Spearman rank correlation $\rho$ between GC% and mutation biases$\Delta M$ as in (A) for all codons. All correlations were statistically significant ("*", $p < 0.05$) after correcting for multiple hypothesis testing. Note that "Z" indicates the serine codons AGC/AGT, which are treated as separate from the other 4 serine codons.

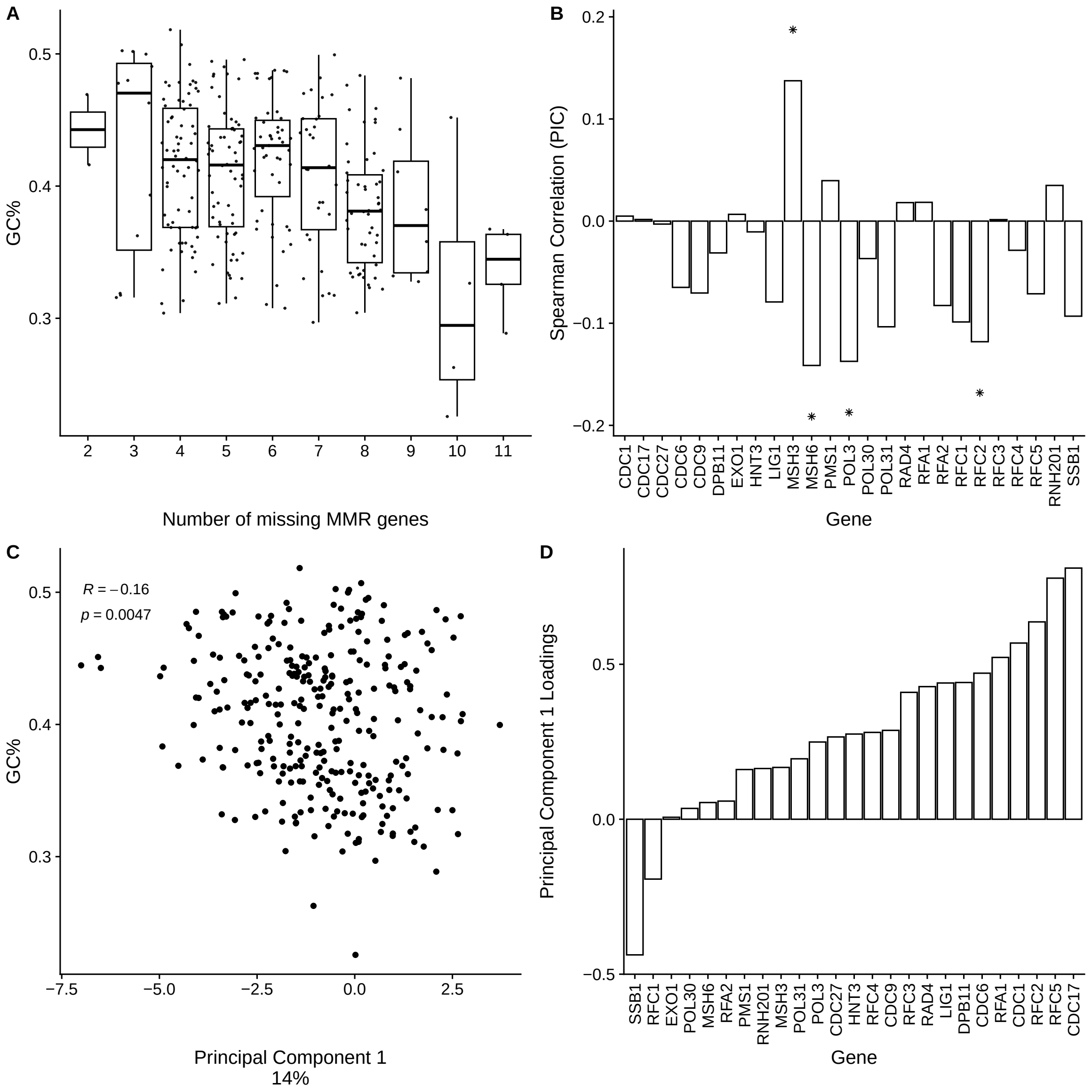


Figure S23: Relationship between mismatch-repair (MMR) gene expression and GC%. (A) GC% as a function of the number of missing MMR genes based on (14). A PGLS based on these two groups did not reveal a significant difference in the mean GC% ($\beta = -0.0027, p = 0.5003$). (B) Spearman rank correlations between ROC-SEMPPR estimates of gene expression $\phi$ and GC% across species for various MMR genes. Phylogenetic-independent contrasts were applied in all cases. "*" indicates statistical significance $p < 0.05$ after correcting for multiple hypothesis testing. (C) Correlation between GC% and the first principal component scores based on a phylogenetic-PCA applied to the predicted gene expression values $\phi$ of 25 MMR (D) Variable loadings from phylogenetic-PCA.

### References


[1] Shen XX, Opulente DA, Kominek J, Zhou X, Steenwyk JL, Buh KV, et al. Tempo and Mode of Genome Evolution in the Budding Yeast Subphylum. Cell. 2018 11;175:1533-45.e20.

[2] Revell LJ, Harmon LJ, Collar DC. Phylogenetic Signal, Evolutionary Process, and Rate. Systematic Biology. 2008 8;57:591-601. Available from: https://dx.doi.org/10.1080/10635150802302427.

[3] Hansen TF. Stabilizing selection and the comparative analysis of adaptation. Evolution. 1997 10;51:1341-51. Available from: http://doi.wiley.com/10.1111/j.1558-5646.1997.tb01457.x.

[4] Pennell MW, Eastman JM, Slater GJ, Brown JW, Uyeda JC, Fitzjohn RG, et al. geiger v2.0: an expanded suite of methods for fitting macroevolutionary models to phylogenetic trees. Bioinformatics. 2014 8;30:2216-8. Available from: https://dx.doi.org/10.1093/bioinformatics/btu181.

[5] Rogalski M, Karcher D, Bock R. Superwobbling facilitates translation with reduced tRNA sets. Nature Structural and Molecular Biology. 2008 2;15:192-8.

[6] Rocha EPC. Codon usage bias from the tRNA's point of view: Redundancy, specialization, and efficient decoding from translation optimization. Genome Reasearch. 2004;14:2279-86.

[7] Sharp PM, Bailes E, Grocock RJ, Peden JF, Sockett RE. Variation in the strength of selected codon usage bias among bacteria. Nucleic Acids Research. 2005;33:1141-53.

[8] Subramanian S. Nearly Neutrality and the Evolution of Codon Usage Bias in Eukaryotic Genomes. Genetics. 2008 4;178:2429-32. Available from: https://dx.doi.org/10.1534/genetics.107.086405.

[9] Vieira-Silva S, Rocha EPC. The Systemic Imprint of Growth and Its Uses in Ecological (Meta)Genomics. PLoS Genetics. 2010;6:1-15.

[10] Labella AL, Opulente DA, Steenwyk JL, Hittinger CT, Rokas A. Variation and selection on codon usage bias across an entire subphylum. PLoS Genetics. 2019 7;15:e1008304. Available from: https://doi.org/10.1371/journal.pgen.1008304.

[11] dos Reis M, Savva R, Wernisch L. Solving the riddle of codon usage preferences: a test for translational selection. Nucleic Acids Research. 2004;32:5036-44.

[12] Hershberg R, Petrov DA. Evidence That Mutation Is Universally Biased towards AT in Bacteria. PLoS Genetics. 2010 9;6:e1001115. Available from: https://dx.plos.org/10.1371/journal.pgen.1001115.

[13] Zheng Q, Steenwyk JL, Rokas A. Lack of universal mutational biases in a fungal phylum. bioRxiv. 2022 3:2022.03.29.486229. Available from: https://www.biorxiv.org/content/10.1101/2022.03.29.486229v2 https://www.biorxiv.org/content/10.1101/2022.03.29.486229v2.abstract.

[14] Phillips MA, Steenwyk JL, Shen XX, Rokas A. Examination of Gene Loss in the DNA Mismatch Repair Pathway and Its Mutational Consequences in a Fungal Phylum. Genome Biology and Evolution. 2021;13.
